# Cap-dependent translation initiation monitored in living cells

**DOI:** 10.1101/2021.05.21.445166

**Authors:** Valentina Gandin, Brian P. English, Melanie Freeman, Louis-Philippe Leroux, Stephan Preibisch, Deepika Walpita, Maritza Jaramillo, Robert H. Singer

**Affiliations:** Janelia Research Campus, Howard Hughes Medical Institute, Ashburn, Virginia, USA; Institut National de la Recherche Scientifique (INRS)-Centre Armand-Frappier Santé Biotechnologie (CAFSB), Laval, Quebec, Canada

## Abstract

Despite extensive biochemical, genetic, and structural studies, a complete understanding of mRNA translation initiation is still lacking. Imaging methodologies able to resolve the binding dynamics of translation factors at single-cell and single-molecule resolution are necessary to fully elucidate regulation of this paramount process. We fused tags suitable for live imaging to eIF4E, eIF4G1 and 4E-BP1 without affecting their function. We combined Fluorescence Correlation Spectroscopy (FCS) and Single-Particle Tracking (SPT) to interrogate the binding dynamics of initiation factors to the 5’cap. Both FCS and SPT were able to detect eIF4E:eIF4G1 binding to the mRNA in the cytoplasm of proliferating cells and neuronal processes. Upon inhibition of phosphorylation by mTOR, 4E-BP1:eIF4E complexes rapidly dissociated from the 5’cap followed by eIF4G1 dissociation. Imaging of the binding dynamics of individual translation factors in living cells revealed the temporal regulation of translation at unprecedented resolution.

## Introduction

Tight regulation of mRNA translation is required to maintain proper cell physiology. Aberrant mRNA translation results in a broad spectrum of diseases, including cancer^1^, metabolic and neurodegenerative disorders^2^, as well as viral infection^3^. Therefore, a better understanding of such (dys)-regulation is key to identify new therapeutic targets.

Single-molecule detection of mRNAs revealed differential distributions in subcellular compartments in neurons, fibroblasts and epithelial cells fueling the hypotheses that translation should be locally regulated to selectively synthesize the proteins where needed^4^. More recently, optogenetic control of β-actin mRNA translation has demonstrated that activation of its translation at the leading edge is required for cell migration^5^. Regulation of mRNA translation is very heterogenous, with mRNAs being differentially translated based on their structure, post-transcriptional modifications and mRNA sequence^6^. It remains unclear which role the diversity of translation factors and ribosomes play in fine-tuning translation of individual mRNAs in subcellular compartments^7^.

Imaging methodologies capable of resolving the regulation of mRNA translation with sub-cellular and single-molecule resolution are necessary to address the impact of local translation in cell physiology. Translational regulation presents a challenge for any single imaging methodology due to its broad range in concentration (from a single mRNA to hundreds of thousands of ribosomes and translation factors per cell), as well as the broad temporal dynamic range (from microseconds binding/unbinding of translation factors, to minutes-long translation times).

In order to address this, we used complementary approaches: single particle tracking (SPT) and fluorescence correlation spectroscopy (FCS). FCS provides ensemble-averaged dynamics of thousands of molecules in seconds, localized to a diffraction-limited volume within the cell. SPT gives a detailed cellular overview of molecular diffusion and utilizes the power of tracking single molecules, but their trajectories have to be recorded individually, which requires low molecular density. The complementarity of the two techniques extends to their temporal dynamic ranges. FCS can detect molecular binding/unbinding events with microsecond temporal resolution, but dynamics are limited to diffusion times of molecules though the focal volume of the laser excitation spot. The temporal resolution of SPT is limited to milliseconds by the camera read-out speed, but may extend over many minutes, ultimately being only limited by the photon budget of the fluorophore of choice. Here we combine these approaches into a powerful methodology capable of resolving molecular interactions of translation factors with mRNA in living cells.

Initiation of cap-dependent translation is considered a rate-limiting step in translation. The best characterized regulatory step is the assembly of eIF4F complexes on the 5’cap structure of the mRNA, which preferentially stimulates translation of a subset of mRNAs that control cell growth, proliferation, metabolism and cell survival^8^. eIF4F assembly is regulated by the mechanistic target of rapamycin (mTOR). At initiation, mTOR phosphorylates a family of inhibitory proteins called 4E-BPs^9^. Upon phosphorylation, 4E-BPs are released from eIF4E exposing the eIF4G binding site^10^. This allows eIF4G to bind eIF4E to facilitate recruitment of other initiation factors to stabilize the small ribosomal subunit on the mRNA^8^. We monitored binding of eIF4F complexes to the mRNA as a proof-of-principle by labeling individual molecules of the initiation factors eIF4E1 and eIF4G1, and then detecting their changes in diffusion by FCS and SPT when they bind to the mRNA. Using the same methodology, we investigated the diffusional behavior of the translation repressor 4E-BP1 and its effect on eIF4E. We were able to detect hotspots of local translation initiation in the cytoplasm of dividing cells as well in neuronal processes, with single mRNA resolution.

## Results

### Dynamic detection of eIF4E bound to the 5’cap of mRNAs

FCS reveals the timescale of fluctuations as fluorescent molecules diffuse through the focal volume in crowded conditions. Therefore, we reasoned that FCS had the potential to resolve differential diffusion of initiation factors, mRNA-bound vs -unbound, despite their high concentration in living cells. The focal volume is in the order of femtoliters with detection limits ranging from 100pmol to 100nmol. Since diffusion of thousands of molecules are averaged in tens of seconds, autocorrelations determine diffusion of molecules with excellent statistical confidence in a short time, presumably without locally perturbing mRNA translation.

The Halo-tag was fused to the non-conserved N-terminal region of eIF4E that is dispensable for eIF4G binding^11^. Since eIF4E is an essential gene^12^, cells where it is knocked down stop proliferating. Halo-eIF4E was made insensitive to eIF4E shRNA by synonymous changes in the coding sequence and was expressed in NIH3T3 fibroblasts in which the endogenous counterpart was silenced by shRNA (Suppl. Fig 1a). Exogenous Halo-eIF4E was able to bind the 5’cap and rescue cell proliferation in NIH3T3 cells (Suppl. Fig 1b,c). These data demonstrated that the Halo tag did not perturb eIF4E function. We then determined whether FCS could resolve _JF-646_Halo-eIF4E binding to the 5’cap in living NIH3T3. Prolonged slow fluctuations in the fluorescent signal, corresponding to slow moving molecules throughout the focal volume, were observed in the cytoplasm, but not in the nucleus where translation does not occur (Suppl. Fig 1d). Autocorrelations confirmed that two-eIF4E components exist in the cytoplasm of translating cells (“faster” and “slower”). Only one fast-component is detected in the cytoplasm or nucleus (Fig 1a right, Suppl. Fig. 1e) after a two-hour treatment with the mTOR inhibitor torin-1, which promotes dephosphorylation of 4E-BP1 whereupon it binds to eIF4E and inhibits eIF4E binding to the 5’ cap, thus preventing initiation. This inhibition was confirmed by a decrease in the expression of Cyclin D1, a canonical eIF4E-sensitive mRNA^13^ (Fig. 1b).

**Fig. 1.**
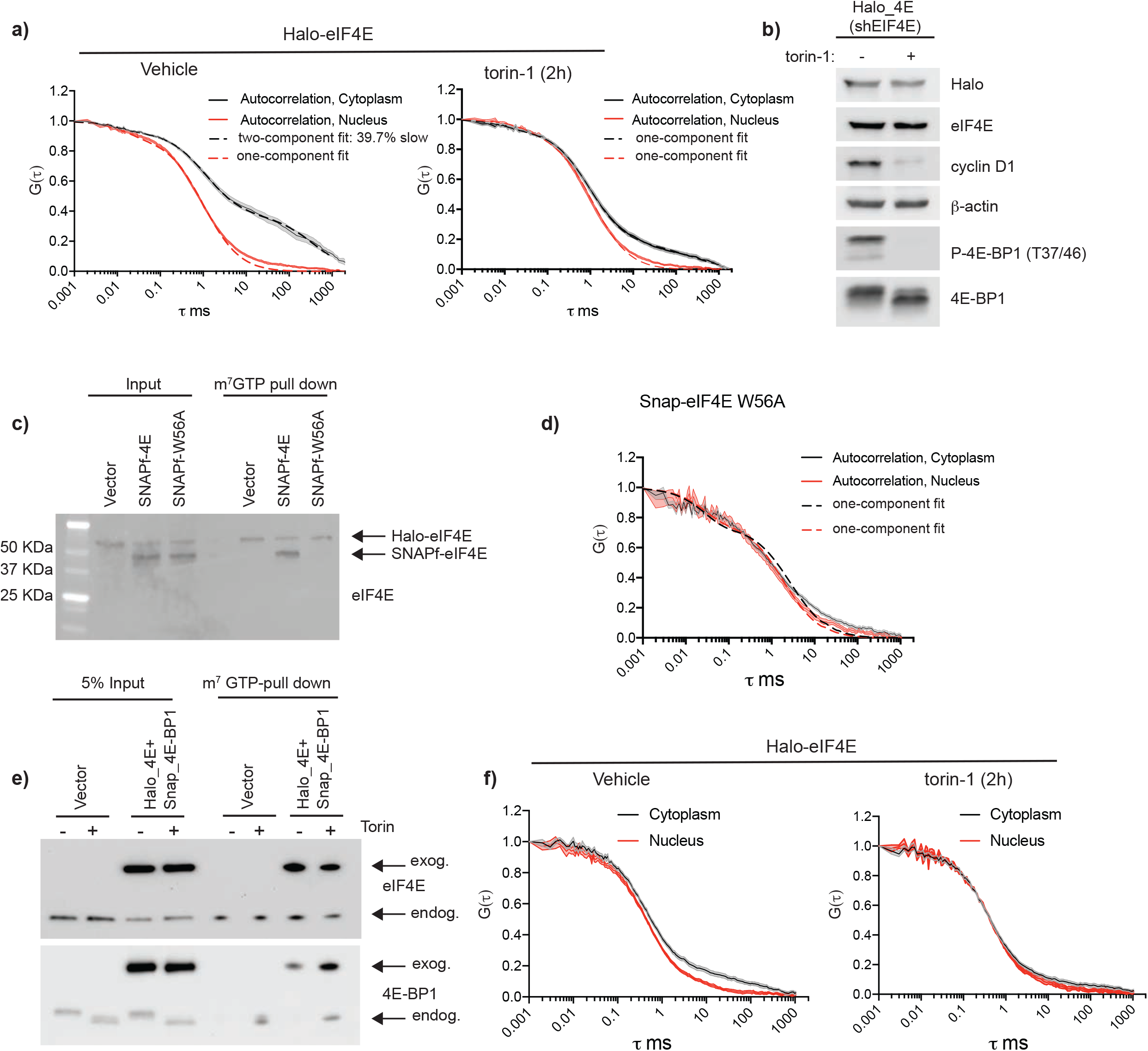
Binding of exogenous Halo-eIF4E to the 5’cap is detected by FCS. **a)** NIH3T3 cells that express Halo-eIF4E, in which the endogenous counterpart was silenced by shRNA, were treated for 2h with vehicle (DMSO) or 250nM torin-1. Autocorrelation curves show diffusion speed of Halo-eIF4E molecules (ms=milliseconds) in the cytoplasm (black) and in the nucleus (red) in the indicated conditions. Halo-eIF4E diffusion is slower in translating cells as compared to cells treated with torin-1. Cytoplasmic autocorrelation best fit with two components (fast and slow). Percentage of slow-moving molecules is indicated. **b)** Total cell lysates from the cells described in (a) were analyzed by western blotting with the indicated antibodies. 4E-BPs phosphorylation and CyclinD1 expression were significantly reduced after 2h torin-1. **c**,**d)** SNAPf-eIF4E and SNAPf-eIF4E^W56A^ (SNAPf-W56A) were expressed in the cells described above. Total cell lysates were subjected to cap-pull down assay. Levels of Halo-4E, SNAPf-4E and SNAPf-4E W56A were detected by western blotting in input (5%) and cap-bound fractions (c) using eIF4E antibodies (eIF4E). Autocorrelation curves show diffusion speed of SNAPf-eIF4E W56A in the cytoplasm (red) and in the nucleus (black). Only one fast component was detected in both cellular compartments (d). **e**,**f)** NIH3T3 cells that express both Halo-eIF4E and SNAPf-4E-BP1 were treated with vehicle (DMSO) or 250nM torin-1 for 2h. Total cell lysates were subjected to cap-pull down assay and analyzed by western blotting in the indicated fractions Overexpression of SNAPf-4E-BP1 is sufficient to increase its binding to eIF4E on the 5’cap (e). Autocorrelation curves show diffusion speed of Halo-4E in the indicated conditions. SNAPf-4E-BP1 expression is sufficient to displace most of the Halo-4E bound to the 5’cap (f).

In order to verify that the slower component of eIF4E diffusion in the autocorrelation was due to its binding to the 5’cap of the mRNA, the eIF4E^W56A^ mutant was substituted for the wild type. Changing tryptophan 56 to alanine perturbs the electrostatic interactions to the 7-methyl-guanine of the 5’cap preventing eIF4E from binding to the mRNA^14,15^. The shRNA-resistant Halo-eIF4E and either the SNAP_f_-eIF4E or the SNAP_f_-eIF4E^W56A^ were co-expressed in NIH3T3 in which their endogenous counterpart had been silenced by shRNA. As expected, the wild-type version of SNAP-tagged eIF4E was able to bind the 5’cap, while the Trp56Ala mutation abolished the binding to the 5’cap (Fig. 1c). Autocorrelations of cytoplasmic and nuclear SNAP_f_-eIF4E^W56A^ revealed only one-fast component in translating cells (Fig. 1d), consistent with the conclusion that the slow diffusion of eIF4E in translating cells was due to its binding to the mRNA.

The eIF4F complex formation is inhibited in the cytoplasm by 4E-BP binding to eIF4E upon mTOR inhibition^9^. We next sought to determine whether the dissociation of eIF4E from the 5’cap was triggered by 4E-BP1 binding. SNAP_f_ tag was fused to the N-terminus of 4E-BP1, in order to preserve the TOR signaling (TOS) motif at the C-terminus. 4E-BP1/eIF4E stoichiometry determines the sensitivity to active-site mTOR inhibitors (asTOR_i_), which are more efficient with higher 4E-BPs levels ^16^. To assess whether the SNAP_f_ tag perturbed 4E-BP1 function, a shRNA-resistant SNAPf-4E-BP1 was expressed in NIH3T3 in which the endogenous counterpart was depleted by shRNA (Suppl. Fig. 2a). As expected, 4E-BP1 overexpression attenuated cell proliferation compared to the vector control, with or without the endogenous knockdown (Suppl. Fig. 2b). NIH3T3 cells that express vector control or SNAP_f_-4E-BP1 were treated with a vehicle control or torin-1 for 16 hours (Suppl. Fig. 2c). As expected, the cytostatic effect of torin-1 was more pronounced in NIH3T3 that express SNAPf-4E-BP1, due to its enhancement of cap binding and consequent competition with eIF4G. These data demonstrate that the SNAP_f_ tag does not perturb 4E-BP1 function. In vitro cap-binding assays confirmed that overexpression of 4E-BP1 was sufficient to increase its binding to eIF4E, with a more pronounced interaction upon mTOR inhibition (Fig. 1e). In living cells, FCS revealed that 4E-BP1 overexpression correlated with _JF646_Halo-eIF4E dissociation from the 5’cap, that was further increased upon mTOR inhibition (Fig. 1f).

**Fig. 2.**
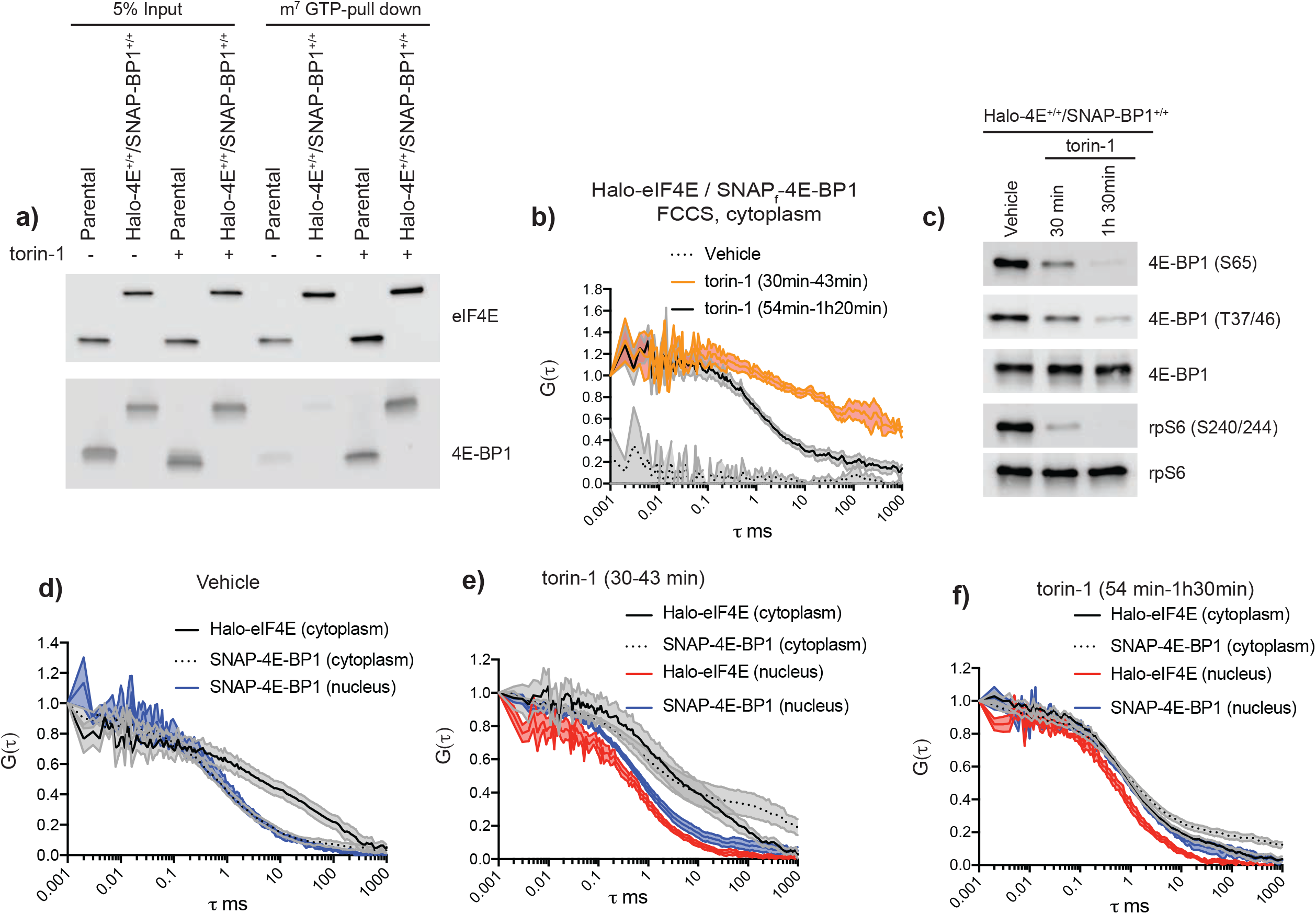
eIF4E is released from the 5’cap upon binding to 4E-BP1. **a)** Differentiated parental and mESC in which Halo and SNAP_f_ tag were inserted into the EIF4E1 and 4EBP1 locus respectively (Halo-4E^+/+^/SNAP-BP1^+/+^), were treated with vehicle (DMSO) or 250nM torin-1 for 1hour and 30 minutes. Total cell lysates (Input) were subjected to cap-pull down assay and analyzed by western blotting using the indicated antibodies. The Halo and SNAP_f_ tags do not affect eIF4E:4E-BP1 binding upon mTOR inhibition. **b)** mESC double knock-in described in (a) were treated with vehicle (DMSO) or 250nM torin-1. Simultaneous diffusion of _JF585_Halo-eIF4E and _JF646_SNAPf-4E-BP1 was analyzed by dual color cross-correlation spectroscopy in the indicated conditions. Cross-correlation was detected, in the cytoplasm, 30 to 1h 20 minutes upon mTOR inhibition and with differential diffusion speed. The vehicle showed no correlation over time (gray). **c)** mESC double knock-in described in (a) were treated with vehicle (DMSO) or 250 nM torin-1 for 30 or 90 minutes. Total cell lysates were analyzed by western blotting using the indicated antibodies. 4E-BP1 and rpS6 were used as loading controls. **(d-f)** Individual diffusion of _JF585_Halo-eIF4E1 and _JF646_SNAP_f_-4E-BP1 was analyzed by FCS in control cells (vehicle) (d) or in torin-1 treated cells (e,f). Autocorrelation curves show two Halo-eIF4E1 components (fast and slow) and one-fast Snapf-4E-BP1 component in the cytoplasm of translating cells. Nuclear diffusion of SNAP_f_-4E-BP1 is depicted in blue (d). Upon 30-43 minutes torin-1 treatment, SNAP_f_-4E-BP1 diffusion slows down in the cytoplasm with Halo-eIF4E still moving slower than its nuclear counterpart (e). After 54-90 minutes torin-1 treatment, both Halo-eIF4E and SNAP_f_-4E-BP1 autocorrelations show overall fast diffusion (f).

To avoid experimental variations in eIF4E:4E-BP1 stoichiometry that affect the dynamics of translation regulation, Halo and SNAP_f_ tags were inserted into the endogenous EIF4E1 and EI4EBP1 loci respectively using Crispr/Cas9 in mouse Embryonic Stem Cells (mESC) (Fig. 2a). Since cap-dependent translation is more pronounced throughout mESC differentiation^17^, parental and double knock-in (DKI) Halo-eIF4E^+/+^/SNAP_f_-4E-BP1^+/+^ mESC were differentiated into fibroblasts (not shown). mTOR inhibition promoted eIF4E binding to 4E-BP1 in both parental and DKI with no major difference (Fig. 2a). We then examined the diffusional behavior of _JF646_SNAP_f_-4E-BP1 and _JF585_Halo-eIF4E by Fluorescence Cross-Correlation Spectroscopy (FCCS). FCCS detected the simultaneous diffusion of the two fluorescent tags (Fig. 2b) through the focal volume for thousands of molecules. The photon counts as a function of time for the two fluorescent channels were correlated with each other, the amplitude of which reflected their simultaneous occupancy in the same complex^18,19^. The cross-correlation analysis demonstrated almost no correlation in translating cells, indicating that the tagged molecules were diffusing freely (Fig. 2b, vehicle). After 30 minutes of mTOR inhibition, partial 4E-BP1 dephosphorylation was observed (Fig. 2c) with _JF646_SNAP_f_-4E-BP1 and _JF585_Halo-eIF4E molecules interacting together with the mRNA as demonstrated by the slow component in the cross-correlations (Fig. 2b). After 1 hour, mTOR inhibition led to a more pronounced 4E-BP1 dephosphorylation (Fig 2c) with the eIF4E:4E-BP1 slow component of the complexes (representing co-binding to the mRNA) almost eliminated (Fig. 2b). These data indicate that 4E-BP1 initially bound eIF4E at the 5’cap to trigger its subsequent dissociation from the mRNA. Autocorrelations for single _JF646_SNAP_f_-4E-BP1 and _JF585_Halo-eIF4E signals demonstrated that 4E-BP1 diffusion significantly slowed down in the cytoplasm after 30-minutes with simultaneous changes in the diffusion of eIF4E molecules compared to control cells (Fig. 2d,e). As expected, mTOR inhibition did not perturb 4E-BP1 and eIF4E diffusion in the nucleus, suggesting that the binding events occurred on the mRNA to inhibit translation initiation. After 1 hour, FCS revealed that both 4E-BP1 and eIF4E diffuse faster than at 30 minutes (Fig. 2f), most likely due to the release of the eIF4E:4E-BP1 complexes from the mRNA. Within 2-3 hours of torin-1 treatment, residual eIF4E binding to the mRNA was not detected, as demonstrated by the cytoplasmic autocorrelations overlapping with the nuclear autocorrelations (Suppl. 3a). After 2-3 hours, eIF4E molecules translocate to the nucleus in a complex with 4E-BP1 as demonstrated by FCCS and fluorescent microscopy. An enrichment in 4E-BP1 molecules was observed in the cytoplasm, most likely to prevent translation initiation, since eIF4E:4E-BP1 complexes are still detected by FCCS (Suppl. 3b,c).

### Dynamics of eIF4E1:eIF4G1 binding to the mRNA

FCS and FCCS were able to detect, in real-time, binding and unbinding dynamics of eIF4E to the 5’cap mediated by the inhibitory molecule 4E-BP1. We reasoned that, if these events occurred on the mRNA, FCS should be capable of detecting changes in the dynamics of the eIF4F complex binding upon acute mTOR inhibition. Halo and SNAP_f_ tags had been inserted into the eIF4E and eIF4G loci respectively at the N-terminus as described above. The SNAP_f_ tag was fused to the N-terminus of eIF4G to preserve the C-terminus which contains the MAPK-interacting kinases 1 (Mnk-1) binding site^20^. Mnk1 is recruited on the eIF4F complex by eIF4G to phosphorylate eIF4E at Serine 209. Since eIF4E phosphorylation stimulates translation of specific mRNAs^21^, we did not want to perturb this regulatory step. Tagging of endogenous eIF4E and eIF4G did not perturb their binding to the 5’cap as compared to the endogenous counterparts lacking the Halo and SNAP_f_ tags respectively (Fig. 3a). We then assessed the simultaneous diffusion of _JF646_SNAP_f_-eIF4G and _JF585_Halo-eIF4E1 by FCCS in the cytoplasm and in the nucleus. As expected, FCCS detected stable eIF4E:eIF4G interactions in the cytoplasm, where mRNA translation occurs, but not in the nucleus (Fig. 3b, left panel). More importantly, their interactions were perturbed in the cytoplasm upon mTOR inhibition (Fig. 3b, right panel). Autocorrelation analysis revealed further insights into their binding dynamics. In control cells, both eIF4E and eIF4G cytoplasmic autocorrelations show that they diffuse as two-components with the majority of the eIF4G molecules moving on average slower than eIF4E (Fig. 3c,d left panel). In the nucleus only one-fast component was detected for both molecules (Fig. 3 c,d right panel). Upon two hours of mTOR inhibition, Halo-eIF4E was released from the mRNA (Fig. 3c, left panel), as previously observed in the double knock-in Halo-eIF4E/SNAPf-4E-BP1. This coincided with a drop in the polysome levels accompanied by monosome accumulation, resembling defects in translation initiation (Suppl. Fig. 4a). Unlike eIF4E, major changes in the diffusion of cytoplasmic SNAP_f_-eIF4G were observed only after five hours of mTOR inhibition (Fig. 3d, left panel). After prolonged mTOR inhibition, cytoplasmic eIF4G autocorrelations mirrored the nuclear fast component (Fig. 3d), suggesting that eIF4G may reside longer on the mRNA or in ribonucleoprotein complexes upon the release of eIF4E from the 5’cap. Since eIF4E is the scaffold protein that recruits eIF4G on the mRNA, this result was unexpected. We then asked whether fusion of the SNAP_f_ and Halo tags to eIF4G and eIF4E respectively could impair eIF4F complex binding dynamics. In vitro cap-pull down assays demonstrated that tagged eIF4E:eIF4G interactions remain mTOR-dependent (Fig.3e), suggesting that the unexpected kinetics observed in living cells are not caused by the presence of the genetically encoded tags. To further rule out the possibility that tagging of two initiation factors alters their binding dynamics in live cells, we performed FCS in Halo-eIF4E or SNAP_f_-eIF4G knock-in differentiated mESCs (Suppl. Fig. 5a-c). The same binding dynamics were observed in these cells, with eIF4G also bound longer to the mRNAs as compared to eIF4E upon mTOR inhibition (Suppl. Fig 5d,e). In order to investigate whether these unexpected binding dynamics were unique to differentiated mESCs, SNAP_f_-eIF4G and Halo-eIF4E1 were expressed in NIH3T3 cells (Fig. 4a). Dual color FCCS was able to detect stable eIF4E:eIF4G interactions in the cytoplasm, but not in the nucleus (Fig.4b, left panel). Acute mTOR inhibition abolished most of the eIF4E:eIF4G interactions as previously observed in differentiated mESCs (Fig. 4b, right panel). Individual autocorrelations confirmed that eIF4E, unlike eIF4G, was rapidly released from the mRNA (Fig. 4c,d) to retard translation initiation as was observed by polysome profiling (Suppl. Fig 4b).

**Fig. 3.**
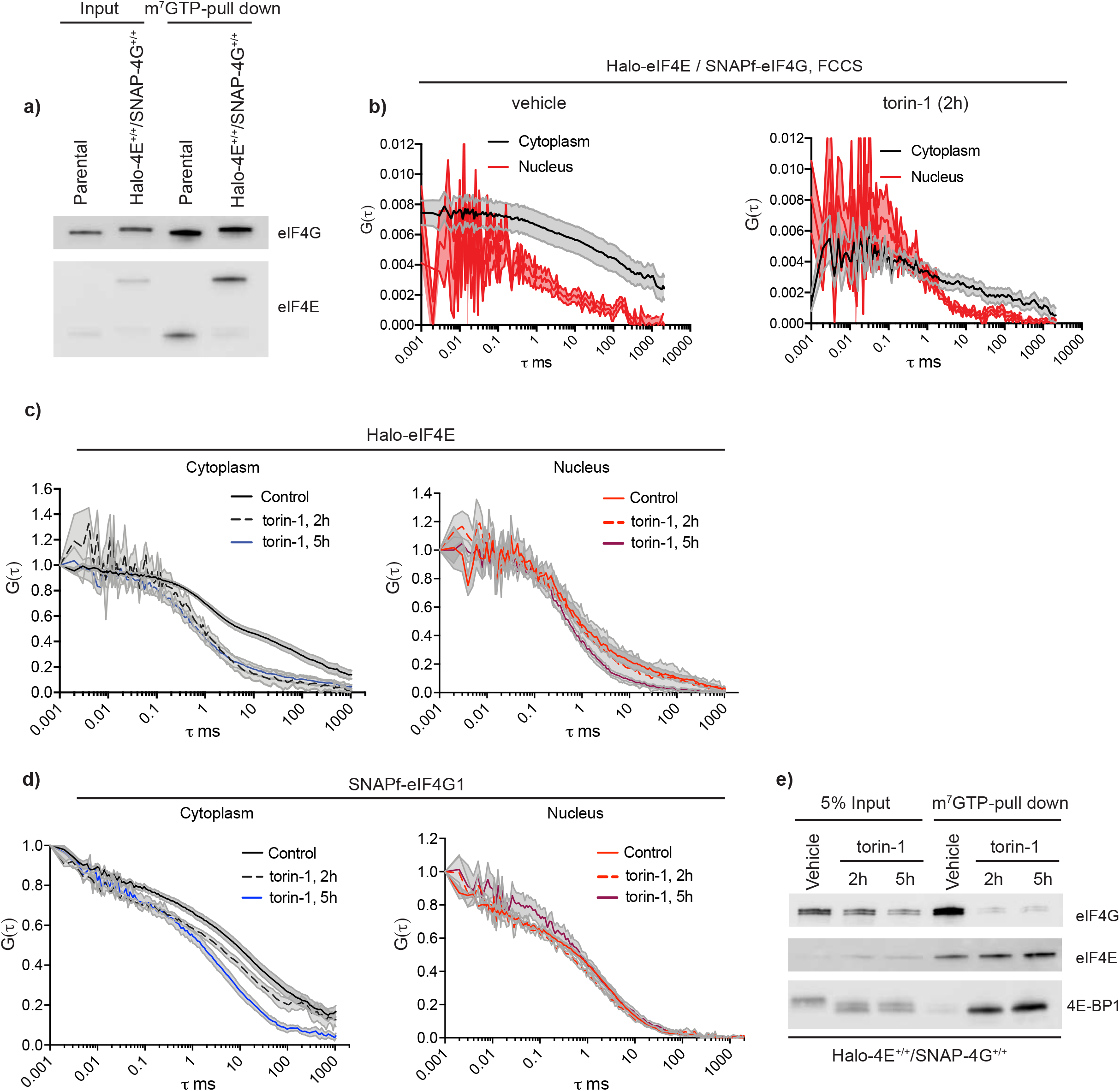
eIF4E and eIF4G binding to the mRNA is detected by FCS upon mTOR inhibition. **a)** Parental mESC and double knock-in Halo-eIF4E^+/+^/SNAP-eIF4G1^+/+^ total cell lysates (input) were subjected to cap-pull down assay (m7GTP-pull down) and analyzed by western blotting. eIF4G and eIF4E antibodies showed binding of wild-type and tagged proteins to the 5’cap. Tagging of endogenous eIF4E and eIF4G did not affect cap-binding as compared to the parental counterpart. **b-e)** Halo-eF4E1^+/+^/SNAP-eIF4G1^+/+^ cells were treated with vehicle (DMSO) or 250nM torin-1 for 2 hours and 5 hours respectively. **b)**Simultaneous diffusion of _JF585_Halo-eIF4E and _JF646_SNAPf-eIF4G1 was analyzed by dual color cross-correlation spectroscopy (FCCS) in the indicated conditions. Cross-correlation was detected in the cytoplasm of control cells, but not in the nucleus (left panel). Minor residual eIF4E:eIF4G was detected in the cytoplasm upon 2 hours mTOR inhibition (right panel). **c**,**d)** Autocorrelation curves representing individual diffusion of _JF585_Halo-eIF4E (c) and _JF646_SNAPf-eIF4G1 (d) in the indicated conditions. In control cells, both _JF585_Halo-eIF4E (c) and _JF646_SNAPf-eIF4G1 (d) autocorrelations showed slower diffusion as compared to their nuclear counterparts. No changes were detected in the nuclear counterparts. Cytoplasmic eIF4E molecules diffuse as fast as the nuclear counterpart as early as 2 hours upon torin-1 treatment, whereas cytoplasmic eIF4G mirrors the nuclear diffusion only after 5 hours. **(e)** Total lysates (input) of cells described in (c-e) were subjected to cap-pull down assay (m^7^GTP pull-down) and analyzed by western blotting with the indicated antibodies. eIF4E:eIF4G dissociation occurred as early as 2h upon torin-1 treatment.

**Fig. 4.**
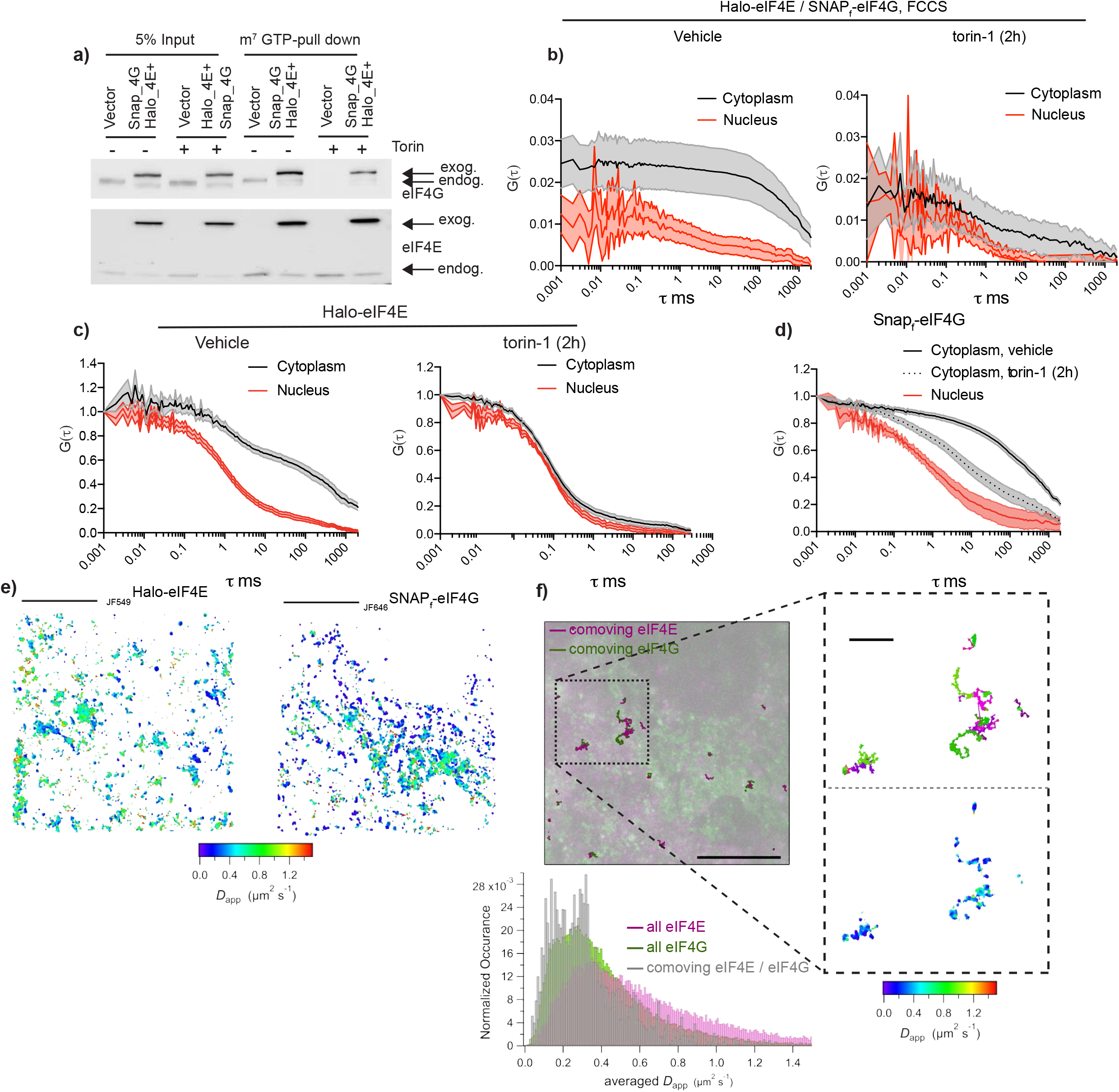
eIF4E and eIF4G binding dynamics detected by FCS and SPT. **a)** NIH3T3 cells that express vector control or Halo-eIF4E and SNAP_f_-eIF4G were treated with DMSO (vehicle) or 250nM torin-1 for 2 hours. Total cell lysates (input) were subjected to cap-pull down assay (m7GTP-pull down) and analyzed by western blotting. eIF4G and eIF4E antibodies detected both endogenous (endog.) and exogenous (exog.) counterpart as indicated by the arrows. **b)** Simultaneous diffusion of _JF585_Halo-eIF4E and _JF646_Snapf-eIF4G analyzed by dual-color fluorescent cross-correlation spectroscopy (FCCS) in cell treated with vehicle (left panel) or 250nM torin-1 for 2 hours (torin-1, right panel) in the cytoplasm (gray) and in the nucleus (red). Cross-correlation was detected in the cytoplasm, but not in the nucleus, of translating cells and abolished 2 hours upon mTOR inhibition. **c**,**d)** Individual _JF585_Halo-eIF4E (c) and _JF646_Snap_f_-eIF4G (d) autocorrelation curves from b). In control cells (vehicle), eIF4E and eIF4G1 molecules diffuse slower than the nuclear counterpart. 2 hours upon torin-1 treatment (torin-1, 2h), eIF4E and eIF4G molecules diffuse faster than in vehicle control. eIF4E, unlike eIF4G, quickly dissociate from the 5’cap upon mTOR inhibition. ***e)*** Simultaneous single-particle tracking of _JF549_Halo-eIF4E and _JF646_Snapf-eIF4G in the cytoplasm of NIH3T3 cells. *Left:* The diffusion properties of _JF549_Halo-eIF4E and _JF646_Snap_f_-eIF4G are displayed via heat maps where _JF646_Snap_f_-eIF4G trends towards slower diffusion (bluer colors). **f)** Co-movement analysis of _JF549_Halo-eIF4E and _JF646_Snapf-eIF4G reveals where the two molecules are interacting. Maximum intensity projections of 5,000 frames of _JF549_Halo-eIF4E (in magenta) and 5,000 frames of _JF646_Snapf-eIF4G (in green) simultaneously acquired at 100Hz. The co-moving _JF549_Halo-eIF4E (bold, magenta) and _JF646_Snap_f_-eIF4G (bold, green) trajectories are displayed on top. Scale bar: 10 μm. Inset top: co-moving _JF549_Halo-eIF4E / _JF646_Snapf-eIF4G trajectories with their associated diffusion heat maps displayed at higher magnification (scale bar: 1 μm). Inset bottom: The distribution of apparent diffusion coefficients shifts to a slower population when _JF549_Halo-eIF4E is comoving with _JF646_Snap_f_-eIF4G (in gray). This implies that the complex is co-moving on mRNAs.

**Fig. 5.**
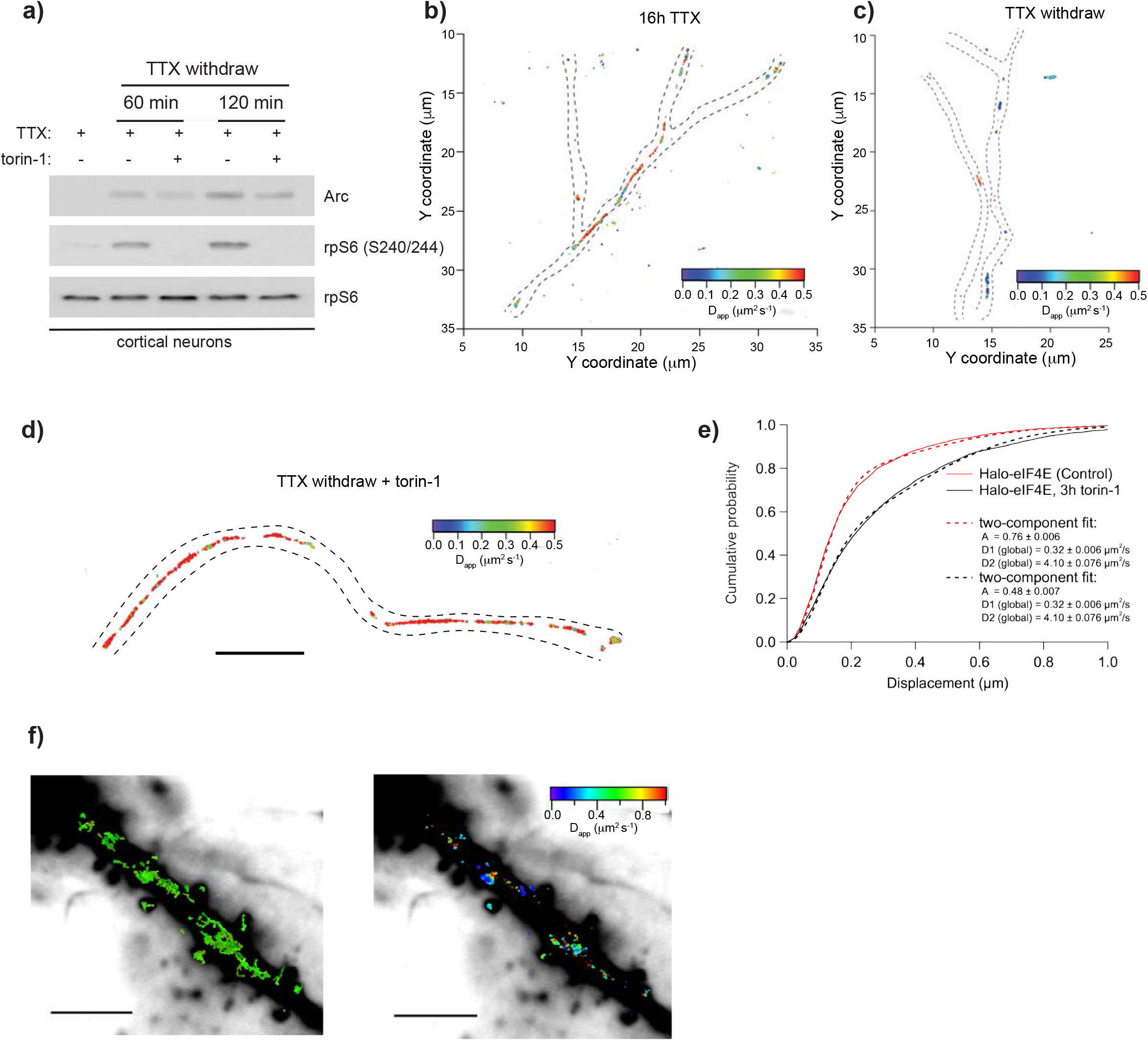
Single-particle tracking of Halo-eIF4E in primary hippocampal neurons. Rat hippocampal neurons were activated with TTX withdraw with or without 250nM torin-1. **a)** Total cell lysates were analyzed by western blotting with the indicated antibodies. TTX and torin-1 inhibit mTOR kinase activity as shown by rpS6 phosphorylation. ARC mRNA indicates the activation. **b)** Diffusion heat map of Halo-eIF4E trajectories obtained in inactivated neurons (16h TTX) at a frame rate of 100 Hz. **c)** Diffusion map of Halo-eIF4E of activated neurons after TTX withdrawal. We detect areas of slow diffusion (in blue). We attribute this to the increased activity of mTOR that leads to the formation of cap-dependent hotspots of translation in dendrites (dashed-outline) of activated neurons. **d)** Inactivating mTOR (with torin-1) these hotspots disappear and the overall diffusion speed of Halo-eIF4E increases (red colors) (scale bar: 5 μm). **e)** The cumulative distribution function of all single molecule displacements for Halo-eIF4E is shifted to the left when compared to neurons treated with torin-1, which indicates a shift towards slower diffusion for Halo-eIF4E undergoing cap-dependent translation. **f)** *Left:* Medium-intensity projection of 10,000 frames of _JF549_Halo-eIF4E recorded simultaneously with sparsely photo-activated _PA-JF646_Halo-eIF4E molecules (individual _PA-JF646_Halo-eIF4E trajectories are shown in green) at 56 Hz frame rate (scale bar: 10 μm). *Right:* The associated _PA-JF646_Halo-eIF4E diffusion heat map depicts that slow Halo-eIF4E molecules (in blue) linger near what appear to be spines in activated neurons (scale bar: 10 μm).

In order to validate these findings with an orthogonal approach, single particle tracking (SPT) was used to investigate the differential diffusion of initiation factors. SPT revealed that eIF4G molecules diffused on average more slowly than eIF4E (Fig. 4 e,f) as had been determined using FCS. The eIF4G apparent diffusion coefficient ranged between 0.1 to 0.8 μm^2^/s. Interestingly, the diffusion coefficient for the free and mRNA-bound large ribosomal subunits is 0.1 and 0.4 μm^2^/s, respectively ^22^. The slow diffusion of eIF4G could be due to association with the small ribosomal subunit through eIF3^23^. Individual trajectories for eIF4E and eIF4G revealed that the eIF4G apparent diffusion coefficient further decreased when co-moving with eIF4E, most likely due to their stable interactions with the mRNA to initiate translation (Fig. 4e histogram). Altogether, these data revealed that both the use of FCS and SPT confirmed the observations regarding the initiation of cap-dependent translation in living cells.

### Real-time detection of cap-dependent translation “hot-spots” in primary neurons

Local mRNA translation has been postulated to play a critical role in neuronal function, including neurite remodeling, synapse formation and pruning, and synaptic plasticity^24^. Upon their transcription, mRNAs are transported into the neuronal processes in a translationally repressed state. During transport, most of the mRNA binding proteins (RBPs) prevent the assembly of the eIF4F complex by blocking eIF4E binding to the 5’cap ^25–27^. As demonstrated above, differential diffusion of initiation factors was a read-out for localized events of cap-dependent translation initiation.

In order to detect cap-dependent translation in neuronal processes, Halo-eIF4E was expressed in primary neurons and its diffusion analyzed upon global neuronal activation by tetrodotoxin (TTX) withdrawal. This method utilized prolonged treatment of cultured neurons with TTX, a sodium channel blocker, followed by its quick washout to trigger quasi-synchronous neuronal activity. TTX withdrawal stimulated mTOR activity within the first two hours (Fig. 5a). Under this activation protocol, a significant portion of eIF4E molecules show slow diffusion in the dendrites (Fig.5c), whereas the majority of the eIF4E molecules diffuse freely along the dendrites in inactivated neurons (Fig 5b) or upon acute mTOR inhibition (Fig.5d). eIF4E movements in activated neurons best fit with two-components: a fast apparent diffusion coefficient of 4-10 μm^2^/s, and a slower one with a coefficient of 0.32 μm^2^/s. Translating mRNAs had an apparent diffusion coefficient of approximately 0.1μm^2^/s ^22^. This suggested that the slow moving eIF4E may reflect early events of translation initiation. Indeed, the two-component fit of eIF4E trajectories upon mTOR inhibition revealed a reduction in the slower diffusion component (from 76% to 48%) (Fig. 5e)

Local mRNA translation at activated synapses is thought to be associated with long-lasting synaptic plasticity^24^. In mature dendrites, we detected clusters of single exogenous Halo-eIF4E molecules lingering near the spines and diffusing slowly at or less than 0.1μm^2^/s (Fig. 5f), suggesting that translation initiation occurred in these areas. In order to avoid artifacts due to overexpression of initiation factors, we then interrogated early events of endogenous eIF4F complex assembly. The double-knock in Halo-eIF4E and SNAP_f_-eIF4G mESC were differentiated into neurons^28^ (Suppl. Fig. 6a). As with primary neurons (Fig. 5a), TTX withdrawal activated mTOR kinase and eIF4E (data not shown). In activated neurons, Halo-eIF4E and SNAP_f_-eIF4G had differential diffusion properties as previously detected by FCS and SPT in fibroblasts (Fig. 6a). Both eIF4E and eIF4G diffusions were best fit with two components, with the faster and slower components having apparent diffusion coefficients of >1μm^2^/s and 0.1 μm^2^/s respectively (Fig. 6b). These data revealed that eIF4F complex formation occurred in dendrites to initiate translation.

**Fig. 6.**
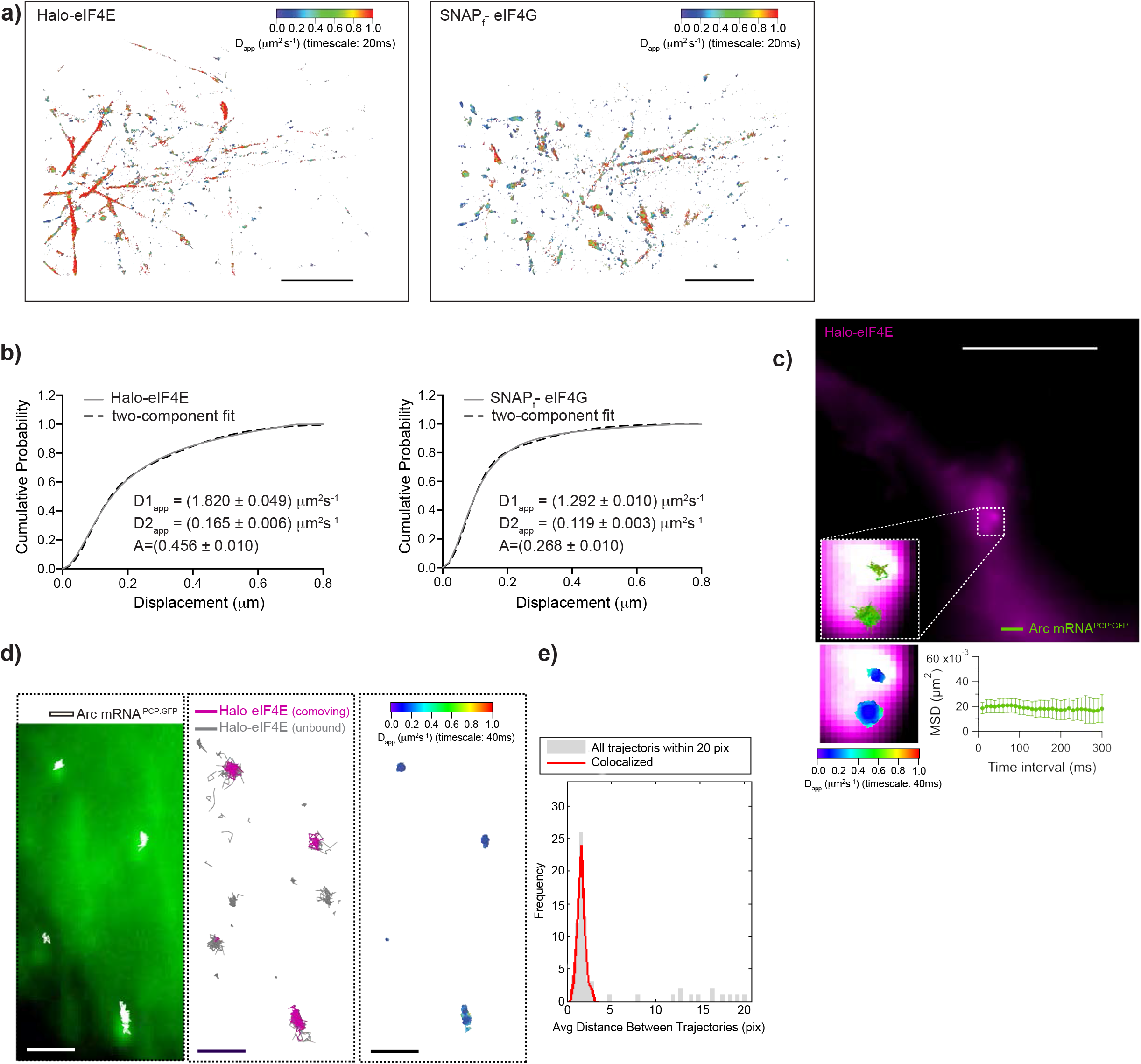
Simultaneous single-particle tracking of endogenous Halo-eIF4E and Snap_f-_eIF4G or Halo-eIF4E and ARC mRNA. **a)** Top left: Diffusion heat map of 18,220 _JF585_Halo-eIF4E trajectories obtained from eight 10,000 frame movies at 50 Hz (scale bar: 10 μm). Top right: Diffusion heat map of the simultaneously recorded 26,438 _JF646_Snapf-eIF4G trajectories. The bluer heat map indicated that eIF4G diffuses more slowly. **b)** The cumulative distribution function, which measures the displacement of the two molecules, shows that Snapf-eIF4G is shifted to the left (slower diffusion) when compared to Halo-eIF4E. **c)** Thick dendritic process of a cortical neuron from ARC^P/+^;PCP-GFP;v-Glut2Cre mice expressing Halo-eIF4E after TTX withdrawal. Inset top: Median projection of the_JF646_ Halo-eIF4E (5,000 frames recorded at 100 Hz). Inset bottom: 2×2 μm^2^ inset of _JF646_ Halo-eIF4E and two ARC mRNA trajectories with their associated diffusion heat maps displayed at higher magnification. The mean square displacement (MSD) curve of the two ARC mRNA trajectories depicts corralling of mRNAs and reveal an exploration area of 0.02 μm^2^ for the 300 ms interval. **d**.Dense area of dendritic branches of cortical neurons from mice described in (c) expressing Halo-eIF4E after TTX withdrawal. Left: Median projection of the PCP-GFP channel (6,000 frames recorded at 100 Hz) outlines a dense region of neuronal processes. Overlaid are the trajectories of four ARC mRNA molecules (in white). Middle: Co-moving Halo-eIF4E molecules (magenta) and unbound Halo-eIF4E molecules (gray) are depicted (scale bar: 2 μm). Right: The diffusion heat map of co-moving mRNA/ Halo-eIF4E molecules depicts almost static movements of the translation factor interacting with ARC-mRNA (scale bar: 2 μm). **e** We generated a matrix of the distances between all detected ARC-mRNA particles and eIF4E particles. The distribution for the entire dataset (gray histogram) consists of a peak of short values (<4 pixels). The peak at short distances is the signature of colocalized mRNA/eIF4E trajectories, while the baseline reflects the concentration of fluorescent eIF4Es non-associated with detected ARC mRNA molecules.

Using SPT, the slow diffusing eIF4E molecules were tested as to whether they indeed bind mRNA. ARC is an immediate early gene rapidly transcribed upon neuronal activation to consolidate long-term memory formation^29^. eIF4E activity has been shown to be rate-limiting for ARC translation^30^ and mTOR inhibition attenuated Arc protein expression without affecting the mRNA stability (Fig. 5a and Suppl. Fig. 6b). In order to label the endogenous ARC mRNA, a triple transgenic mouse was generated that constitutively expressed GFP fused to the coat protein PCP (PCP-GFP). The PCP-GFP tagged the endogenous ARC mRNA in which a tandem array of PP7 binding sites (PBS) had been inserted into the 3’UTR of the ARC gene (Arg 3.1)^31^. Exogenous Halo-eIF4E was expressed in primary neuronal cultures isolated from the ARC-PBS;PCP-2GFP;Vglut2^Cre^ mice. ARC mRNA had been previously shown to be transported along the dendrites, with the anterograde and retrograde transport often interrupted by pauses^31^. However, it remained unclear whether the stationary mRNA was actively translated. In a thick dendrite, a dense area was observed enriched with eIF4E molecules (Fig. 6c). Importantly, two stationary ARC mRNA molecules were detected inside this dense area (Fig. 6c). The binding of eIF4E molecules to the ARC mRNAs was imaged with single-molecule precision. In a dense region of neuronal processes, both fast and slow moving eIF4E molecules were tracked (Fig. 6d). ARC mRNA trajectories were spatially and temporally correlated with eIF4E trajectories to identify a co-moving population for diffusion analysis. Specific co-moving events are apparent when looking at the pooled distances between ARC mRNAs and eIF4E (Fig. 6e). The trajectories were used to generate a diffusion map and demonstrated that mRNA:Halo-eIF4E molecules were almost immobile. This demonstrated that the static translation factors likely represented translation initiation events in neuronal processes.

## Discussion

Several imaging methodologies have been developed in order to visualize the spatial and temporal dynamics of mRNA translation^32–35^. These methodologies rely on a fluorescently labelled mRNA and/or simultaneous labelling of nascent polypeptides from a specific mRNA. To date, methodologies able to elucidate general translation regulation in living cells are lacking. In this work, FCS was able to resolve the diffusion of thousands of molecules in tens of seconds, and when combined with an orthogonal approach using SPT was able to resolve binding dynamics of initiation factors in real-time, regardless of the specific mRNAs being translated. The approach was then applied to interrogate the interactions between the initiation factor eIF4E and a physiologically important mRNA in neurons.

Live imaging demonstrated that, in the cytoplasm, only half of the endogenously labeled cap-binding protein eIF4E is bound to mRNA, with the remaining molecules freely diffusing. Previous work demonstrated that a reduction in eIF4E levels by 50% does not affect mouse development^12^, raising the hypothesis that a surplus of eIF4E may exist. We speculate that the cells benefit from an excess of eIF4E by ensuring homogeneous distribution throughout the cytoplasm to rapidly activate bursts of translation necessary to preserve cellular homeostasis in response to environmental changes leading to, for instance increased mitochondria activity^36,37^ or cell proliferation^38,39^. Live imaging revealed additional insights into the regulatory dynamics of translation. Biochemical approaches, *in vitro*, indicated that 4E-BPs did not affect the cap-binding of eIF4E but blocked interactions between eIF4E and eIF4G^10,40^ thus inhibiting assembly of the eIF4F complexes. In living cells, eIF4E binding to 4E-BP1 triggered the release of eIF4E from the 5’cap within 30 minutes upon mTOR inhibition. This was in contrast to the binding dynamics observed in vitro, in which eIF4E constitutively bound to the 5’cap-analogue. Based on these results, it is possible that in live cells, other factors bind the 5’cap or other regulatory elements of mRNAs in order to displace eIF4E:4E-BP1 complexes to inhibit translation. As an example, La-related protein 1 (LARP1) has been recently described to bind the 5’cap and the adjacent 5’terminal oligopyrimidine (5’TOP) motif thus impeding the binding of eIF4E to the cap on the TOP mRNAs^41^. Since eIF4E regulates a broad category of mRNAs, our findings suggest that similar mechanisms of inhibition may apply to other eIF4E-sensitive mRNAs through cap-binding variants, or by associated RNA-binding proteins yet to be discovered.

Despite being the major cap-binding protein, eIF4E activity is primarily rate-limiting in translating mRNAs characterized by a highly structured 5’UTR^37,42^, a model that has been recently challenged^43–45^. In living cells, mTOR inhibition triggered the release of eIF4E from the mRNA after 30 minutes without affecting global translation as previously reported (Suppl. Fig. 4b,c). Mechanistically, eIF4F assembly (eIF4E, eIF4G and eIF4A) is required to unwind secondary structures and pull the 5’UTR through the mRNA channel until the start codon reaches the P site of the ribosome to initiate the synthesis of the polypeptides^46^. This suggests that mRNAs with short 5’UTRs could be translated in an eIF4F-independent manner, with the ribosome being recruited near the start codon by alternative mechanisms. Interestingly, eIF4E activity is required to translate nuclear-encoded mitochondria mRNAs characterized by a very short 5’UTRs in a eIF4A-independent manner or mRNAs containing Cytosine Enriched Regulator of Translation (CERT) domain in their 5’UTRs^12^. It has been recently proposed that diverse eIF4F complexes may regulate the translation of different classes of mRNAs^46^. It remains an area of investigation whether different eIF isoforms may play a role in fine-tuning the translation of different classes of mRNAs. It is also possible that other factors able to bind the 5’cap can translate a subset of mRNAs in an eIF4E-independent manner in live cells, as recently demonstrated for eIF3^47^ or other factors in different species^48^.

eIF4E is localized in the cytoplasm and in the nucleus, with the nuclear eIF4E immunostaining either enriched in discrete nuclear bodies^49^ or diffuse^50^. In the cell types we have analyzed (NIH3T3 fibroblasts, primary neurons and mESC-derived fibroblasts and neurons), the data support the former. In absence of the spherical bodies, temporal autocorrelations obtained by FCS demonstrated that the majority of the eIF4E molecules inside the nucleus diffuse as one fast-component and are not bound to its target mRNAs as previously observed in cancer cells^51,52^. In agreement, prolonged mTOR inhibition (> 3 hours), leads to the accumulation of endogenous eIF4E:4E-BP1 complexes in the nucleus most likely to prevent the binding of eIF4E to the 5’cap.

This work demonstrated that our approach can identify early-events of cap-dependent translation with sub-cellular and single-molecule resolution. The role of cap-dependent translation in neuronal physiology has been so far demonstrated using genetics and pharmacological approaches^53^, but the role of local translation in neuronal processes remains unclear since these approaches lack sufficient spatial and temporal resolution. By generating a diffusion map of endogenous initiation factors, we were able to detect slow diffusion of eIF4E (<0.1 μm^2^/s) consistent with its binding to the mRNA in dendrites of activated primary and mESC-derived neurons and lingering in the proximity of the spines in mature dendrites. The mechanism behind this slow diffusion of eIF4E was confirmed by its co-movement with ARC mRNA indicating that its binding to the mRNA conferred the slow diffusion properties of the mRNA to the initiation factor. Binding events of endogenous eIF4E:eIF4G were also detected in dendrites of activated neurons. Notably, in neurons as in fibroblasts, we observed an overall differential diffusion with eIF4G diffusing more slowly than eIF4E (1.29 μm^2^/s and 1.82 μm^2^/s respectively). In vitro assays determined that about 80% of eIF4G was bound to ribosomes even in absence of the mRNAs^54^. Only 20% of eIF4E was bound to ribosomes in an eIF4G-dependent manner. This supports a model in which eIF4E recruits a 43S initiation complex already containing eIF4G on the mRNA^55^.

Our approach could be extended to a broader repertoire of translation factors and mRNAs in order to elucidate new mechanisms by which regulation of mRNA translation on differential subsets of mRNAs at precise locations can be affected. This will have important implications in understanding how translation dysregulation leads to pathological conditions.

## Methods

### Cell lines, neuronal cultures and compounds

NIH3T3, obtained from America Type Culture Collection (ATCC) were maintained in phenol red-free Dulbecco’s Modified Eagle Medium supplemented with 10% fetal bovine serum, 1% antibiotic-antimycotic mix (ThermoFisher) and 1% GlutaMAX™ (ThermoFisher). JM8.N4 mouse ESCs from the C57BL/6N (a generous gift from Liu laboratory, Janelia Research Campus) were maintained in Knockout DMEM (GIBCO #10829-018) supplemented with 10% Fetal Bovine Serum ES Cell qualified (ATCC® SCRR-30-2020™), 1X GlutaMAX™ Supplement 100X (GIBCO, #35050-061), 1X MEM Non-Essential Amino Acids Solution, 100X (GIBCO, #11140050), 0.1 mM 2-mercaptoethanol (GIBCO, Cat#21985-023), 1000U/mL LIF (EMD Millipore, # ESG1106), 1μM PD03259010 (Millipore Sigma, #PZ0162), 3μM CHIR99021 (StemCell Technologies, #72052). Medium was renewed every second day. All cell lines were maintained in 5% CO_2_ at 37 °C.

Primary rodent work was conducted according to the Institutional Animal Care and Use Committee (IACUC) guidelines. Newborn pups were euthanized, and cortical tissue dissociated, in papain enzyme (Worthington Biochemicals, ~25 U/cortical pair) in dissection solution (10 mM HEPES pH 7.4 in Hanks’ Balanced Salt Solution) for 30 min at 37°C. Following trituration and filter through a 40 μm strainer, single cells were seeded in Plating media [28 mM glucose (Acros #41095), 2.4 mM NaHCO_3_ (Sigma-Aldrich S-8875), 100 µg/mL transferrin (Calbiochem #616420), 25 µg/mL insulin (Sigma #I-6634 or #I-1882), 2 mM L-glutamine (GIBCO #25030-081), 100 U/mL penicillin, 10 µg/mL streptomycin and 10% fetal bovine serum (heat inactivated; HyClone)] in MEM (GIBCO 51200-038) and NbActiv media (BrainBits, LCC) mixed in a 1:1 ratio. 16 hours later, the media was replaced with Plating media and NbActiv mixed in a 1:33 ratio.

Rat cortical cultures were seeded on Poly-D-Lysine Hydrobromide (PDL) (Sigma) diluted in sterile water at 0.2 ug/ml. Cultures were then maintained by feeding cells twice a week by replacing old media 1:1 with fresh NbActiv media.

Newborn mouse neuronal cultures were plated on PDL and laminin (5μg/mL)-coated plates. Mouse neurons were seeded in Plating media [B27/Neurobasal media, 25uM Glutamax and 5% FBS to mouse Neuron culture media (same as mouse Plating media without FBS in ratio of 1:1. (Gibco #17504044)

Where indicated, cells were treated with 250nM torin-1 (Tocris) dissolved in DMSO or with equal amount of DMSO as a vehicle control. Neuronal cultures were treated with 1.5μM Tetrodotoxin TTX) (Sigma-Aldrich, #554412). After 16 hours, TTX was removed and substituted with conditional media with vehicle control or 250nM torin-1 upon 3 extensive washes with Neuronal Buffer provided by the Janelia Research Campus Media Facility.

### Differentiation of mESC to neurons or fibroblasts

For neuronal differentiation, a single mESC cell suspension was obtained and resuspended in Embryoid body (EB) media [Knockout DMEM (GIBCO, 10829-018), 10% FBS ES Cell Qualified (ATCC® SCRR-30-2020™), 1X GlutaMAX™ (GIBCO, #35050-061), 1X MEM Non-Essential Amino Acids Solution (GIBCO, Cat#11140050) and 55μM 2-mercaptoethanol]. 50 cells/well were plated and EBs grow in AggreWell™400 dish (StemCell Technologies #07010), according to manufacturer’s instructions. Briefly, EBs were maintained for 8 days in EB media in 7% CO_2_ at 37°C. At day 4, EB media was supplemented with 5μM retinoic acid (Sigma, #R2625). Medium was replaced every 48 hours. At day 8, EBs were dissociated and filtered through a cell strainer (Falcon, #352340). Single cell suspension was resuspended in BrainPhys complete media and 1.13×10^5^ cells/cm^2^ were seeded in plates coated with 15μg/mL Poly-L-Ornithine (Sigma #P4957) and 0.5μg/cm^2^ laminin (Gibco #23017-015). Cells were incubated for 2 hours in 7% CO_2_ at 37°C. Media was replaced with fresh BrainPhys complete media after 2, 24 and 48 hours. Brainphys Neuronal Medium (#05790) was supplemented with 2% NeuroCult SM1 neuronal supplement (#05711), 1% N2 Supplement-A (#07152), 20ng/mL BDNF (#78005), 20ng/mL GDNF (#78058), 1mM Dibutyryl-cAMP (#73882), 200nM L-ascorbic acid (Millipore, AX1775-3). All supplements were from Stemcell Technologies unless otherwise specified. Differentiating mESC were maintained in 5% CO_2_ at 37°C. After 48 hours, neurons were fed by replacing half of the media every 5 days.

For fibroblast differentiation, mESC were seeded at low density (~20%) in 10μg/mL fibronectin-coated plates in mESC complete media for 16 hours. mESC were then cultured in absence of LIF and 2i (PD03259010 and CHIR99021) for 48 hours to induce fibroblast differentiation.

### Lentiviral shRNA, Plasmids, Crispr/Cas9 knock-in constructs and generation of cell lines expressing tagged translation factors

Non-targeted shRNA control (Scrambled, SHC216) and the EIF4E shRNA (TRCN0000077477), 4EBP1 shRNA (TRCN0000335449) and EIF4G1 shRNA (TRCN0000096812) targeting the Coding Sequence (CDS) were all from Sigma. Lentiviral backbone pLV-EF1a-IRES-Neo was a gift from Tobias Meyer (Addgene plasmid #85139). Mouse eIF4E, 4E-BP1 and eIF4G1 CDS were synthesized (GeneScript Biotech) by fusing a spacer AGC-GGC-GGA-GGC-GGA-TCC-GGC-GGA-GGC-GGA-AGC (Ser-Gly-Gly-Gly-Gly-Ser-Gly-Gly-Gly-Gly-Ser) at the N-terminal. Each CDS was then subcloned into pLV-EF1a-IRES-Neo. Halo or SNAP tag were then inserted upstream the spacer. In order to generate an shRNA-insensitive CDS, silent mutations were inserted inside the seed-sequence targeted by the shRNA. Viral supernatant for each of the indicated constructs was generated by the Janelia Viral Tool facility. Infection was carried out with 8mg/mL polybrene. 48 hours later, NIH3T3 were selected with 500μg/mL G418 for 7 days, at the end of which, protein expression was analyzed by western blotting. The endogenous counterparts were then silenced by infecting the cells with lentiviral particle carrying shRNA or Scrambled control respectively, as described above. Selection was performed with 5μg/mL puromycin. Protein expression was analyzed by western blotting and the cells maintained in complete DMEM with 500μg/mL G418 + 5μg/mL puromycin. To generate NIH3T3 that expressed both Halo-eIF4E and SNAPf-eIF4G1, cells that expressed SNAPf-eIF4G1 were overlayed with Halo-eIF4E viral supernatant as described above. 48 hours later, cells were stained with 100nM JF_646_-Halo ligand for 30 minutes, washed three times in 1X PBS and sorted.

To generate knock-in mESC, guide RNAs were cloned into pTij-U6-sgRNA-CBh-Cas9-PGK-puroR (provided by the Liu lab, Janelia). Three different gRNAs were designed and experimentally tested for each gene (eIF4E: ENSMUST00000029803.11; EIF4G1: ENSMUST00000115460.7; 4E-BP1 ENSMUST00000033880.6). The following were able to generate homozygote knock-in:

eIF4E #3 - 5’ GAACCGGTGAGTATTGCCTT 3’

eIF4G1 #1 - 5’ GTGCTGGGGGGACCCTAATGTGG 3’

4E-BP1 #11.1 – 5’ GCGTGCAGGAGACATGTCGG 3’

The donor sequence (Halo or SNAPf tag flanked by ~ 700 nucleotides complementary to the targeting sequence) was cloned into pUC19 (provided by the Liu lab, Janelia). 500μg of gRNA and 500μg of donor plasmids were electroporated in 1×10^6^ mESC using P3 Primary Cell 96-well Nucleofector™ Kit (Lonza, PBP3-22500) according to manufacturer’s instruction. After 3 days, mESC were stained with 100nM JF_646_ Halo- or SNAP-tag ligands for 30 minutes and single cells were sorted in a 96-well plate. Clones were expanded, genotyped using the following primers and homozygosity further validated by western blotting.

eIF4E_Genot_F2: 5’ GTGGACCGGGGACTGGGGAGAC 3’

4E-BP1_Genot_F1: 5’ AGTTCTGCCACCGTCATCCCTACC 3’

eIF4G_Genot_F2: 5’ GCCCCGTGGAGCCAGGTTGATA 3’

### Western blotting and antibodies

Western blotting was performed as previously described^56^. Briefly, cells were washed 3 times in ice-cold 1X PBS and scraped in RIPA lysis buffer [10mM Tris HCl pH 8.0, 150mM NaCl, 1% Triton-X100, 0.5% Sodium Deoxycholate, 0.1% Sodium Deoxicholate, 1mM EDTA, 5mM NaF, 10mM β-Glycerophosphate, 1mM Vanadate (New England BioLabs) and cOmplete™, Mini, EDTA-free Protease Inhibitor Cocktail (Millipore)]. Total protein lysates were separated on 12% or 4%-15% Mini-PROTEAN® TGX™ Precast Protein Gels (BioRad), transferred on a nitrocellulose membrane (BioRad) and transferred using a semi-dry apparatus (BioRad). The following antibodies were diluted in [1X Tris Buffered Saline solution (TBS) pH 7.4 (ThermoFisher), 5% bovine serum albumin (Sigma-Aldrich) and 0.1% Tween-20]: anti-eIF4E (monoclonal, BD Transduction Laboratories™ #610269), anti-4E-BP1 (#9644), anti-eIF4G1 (#2498), anti-phosho-4E-BP1 Thr37/46 (#2855), anti-phospho-4E-BP1 Ser65 (#9451), anti-phosho-rpS6 Ser 240/244 (#2215), anti-rpS6 (SantaCruz Biotechnology, #sc-74459) all from Cell Signaling Technology, anti-GAPDH (#G8795) and anti-b actin (#A5441) from Sigma-Aldrich, anti-ARC (SantaCruz Biotechnology, #sc-17839, anti-HaloTag (Promega, #G9211), anti-SNAP-tag (New England Biolabs, #P9310S) and anti-Cyclin D1 (BD Bioscience, #556470). Secondary antibodies, rabbit (Sigma-Aldrich) and mouse (Amersham) IgG HRP linked, were used at 1:5,000. Signals were revealed by Clarity Western ECL Substrate (BioRad) and detected using a ChemiDoc™ MP Imaging System (BioRad, #12003154).

### Imaging system

Stroboscopic single particle tracking was performed using a custom-built three-camera microscope as described in^57^. Briefly, the microscope is equipped with an Olympus 100× NA 1.5 TIRF objective and a custom tube lense (LAO-300.0, Melles Griot), resulting in 133.33x overall magnification. The three Andor iXon Ultra EMCCD cameras (cooled to −80 °C, 17 MHz EM amplifiers, preamp setting 3, Gain 400) were synchronized using a National Instruments DAQ board (NI-DAQ-USB-6363). Lasers (490nm, 561nm, 639nm, all Vortran Stradus lasers) stroboscopically illuminated the sample using peak power densities of ~1.7 kW/cm2 using HiLo illumination. During imaging, cells were maintained at 37 °C and 5% CO2 using a Tokai-hit stage top incubator and objective heater.

### Cellular Labeling Using Halo-tag or SNAP-tag Ligands

Immediately prior to imaging, cells were labeled for single molecule imaging as described in^58^. The labeling conditions for co-tracking in fibroblasts were 15nM SNAP-JF646-SNAP tag ligand^59^ (30min) and JF552-Halo tag ligand (1nM, 15min)^60^. For Halo-eIF4E tracking in neurons we used 100nM JF646-Halo tag ligand (15min), and for co-tracking in neurons we labeled with 100nM JF646-SNAP and JF549-Halo tag ligands (15min). For Halo-eIF4E imaging in spines (see Fig. 4E), we also added PA-JF646-Halo tag ligand (15min, 100nM)^61^. PA-JF646 was photoconverted by 100-μs-long excitation pulses of 407 nm laser light (Vortran Stradus, 50 W/cm2) every second. During the course of image acquisition, the pulse length was increased to 200-μs-long pulses.

### 3-camera in silico registration

Slide-mounted TetraSpeck Fluorescent Microspheres (T14792, Invitrogen) were imaged before and after data acquisition. Short movies (100 frames) of the fluorescent broadband beads in all three channels were registered using the using the descriptor-based series registration (2d/3d+t) Fiji plugin^62^. For more than two-channel bead registration, the frames have to be set as channels (Fiji’s “Re-order Hyperstack” command). Sub-pixel accurate registration was achieved via Gaussian mask localization fitting of spots, which were used to compute affine (2D) transformation models^63^. Since un-biased noise distribution is essential for accurate single particle identification and tracking, the Fiji plugin now allows for nearest-neighbor interpolation that does not change actual pixel intensities and thereby preserves the original noise distribution (as compared to linear or other higher-order interpolation schemes). This update is available via the Fiji Updater (http://fiji.sc/Downloads). It can be found under Plugins > Registration > Descriptor based Registration (2d/3d) and Plugins > Registration > Descriptor based Series Registration (2d/3d+t).

### Single particle tracking and analysis

Trajectories were obtained using DiaTrack (v. 3.04, Semasopht), which identifies and fits the intensity spots of fluorescent particles with 2D Gaussian functions matched to the experimentally determined point-spread function. The diffusion maps were created using tracking routines written in IGOR Pro 8.04 (WaveMetrics) as described in^22^. Briefly, local apparent diffusion coefficients of eIF4E and eIF4G are calculated and mobility is evaluated on a 20 nm × 20 nm x–y grid from the mean square displacements over a timescale of 10 or 20 milliseconds.

We determined comoving trajectories (eIF4E and eIF4G, ARC mRNA and eIF4E) using a previously published analysis^59^. We localized particles and built trajectories in both channels separately. Trajectories that dwelled within 320 nm of one another for at least 10 ms were assigned as colocalized. To assess the colocalization statistics, we generated a matrix of the distances between all detected particles in both channels and computed the corresponding histogram^34^. We then normalized the distance histogram to account for the fact the area covered by each distance bin grows, and plotted the resulting normalized distribution, equivalent to the average density of channel 1 spots observed as a function of distance from channel 2 detections. If trajectories do not colocalize, we would randomly detect channel 1 spots at all positions in the cell without regard for channel 2 positions and therefore we would expect to observe a flat distribution. In the case of colocalization, we would expect an enrichment of short distances corresponding to comoving trajectories.

CDF curves were fit with a two-component fit (equation 1) with *A* (fraction of molecules with the diffusion coefficient *D*_1_), *1-A* (fraction of molecules with the diffusion coefficient *D*_2_), *D*_1_ (diffusion coefficient of the first diffusive species in μm^2^/s), *D*_2_ (diffusion coefficient of the second diffusive species in μm^2^/s), *and t* (frame time in s) as fitting parameters:

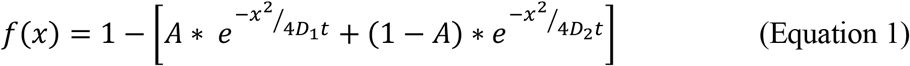

### Fluorescence Correlation Spectroscopy (FCS)

FCS trajectories were obtained using a Leica SP8 Falcon microscope equipped with a Nikon APO 63x 1.20 NA water-immersion objective equipped with a motorized correction collar (mottCorr, Leica). Prior to acquisition, cells at ~70-80% density were labelled with 100nM JF585-Halo tag ligand^64^ and 200nM JF646-SNAP tag ligand^59^ for 10 and 45 minutes, respectively. Cells were heated to 37 °C with 5% CO2 using a Tokai-hit stage top incubator. Just before each measurement, the mottCorr was adjusted using xz-scans in reflection mode to give the sharpest image of the living cell. Xz-scans were was also used to set the central-plane of the cell. A set of two 10s measurements were obtained, one in the nucleus of the cell, and one in the cytoplasm. The FCS curves were exported, and curve-fitting was conducted using DiaTrack (v. 3.04, Semasopht). FCS curves were fitted to a one-component fit (equation 2) with the triplet state fraction of molecules (*T*), the lifetime of the triplet state (*Tt*, in ms), the structural parameter of the focal volume (*κ* = 5), and the diffusion time (*tau*, in ms) as fitting parameters:

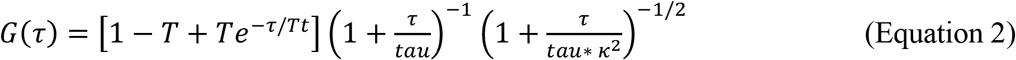

FCS curves were also fitted to a two-component fit (equation 3) with *T* (triplet state fraction of molecules), *Tt* (lifetime of the triplet state in ms), *A* (fraction of molecules with the diffusion time *tau1*), *κ* (structural parameter of the focal volume, *κ = 5*), *tau*_*1*_ (diffusion time of the first diffusive species in ms), and *tau*_*2*_ (diffusion time of the second diffusive species in ms) as parameters:

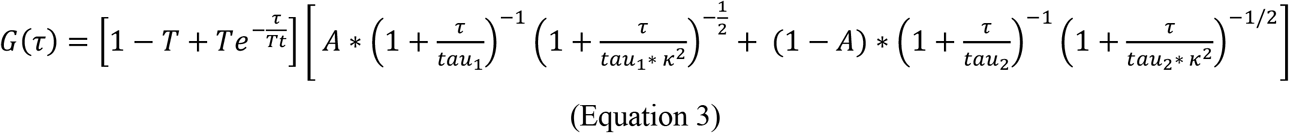

### Cap pull down assay, polysome profiling and ^35^S-Met/Cys labelling

Cap-pull down assay was performed as described in^65^. Briefly, cells at ~70% density were washed 3 times in ice-cold 1X PBS and scraped in Buffer A [50mM Tris HCl pH 7.5, 50mM KCl, 1mM EDTA, 0.5% Sodium Deoxycholate, 1mM Vanadate, 1mM Sodium Fluoride, 20mM b-glycerophosphate]. Total proteins (500μg) were incubated with 15μL Immobilized γ-Aminophenyl-m7GTP (Jena Bioscience, #AC-155) at 4°C for 3 hours. Eluted material was analyzed by western blotting.

Polysome profiling was performed as described in^66^. 5-10 ODs of total RNA, measured at 254 nm, was loaded on a 5%-50% sucrose gradients. Sucrose gradients were displaced with 60% sucrose and the absorbance was recorded continuously at 254 nm using a Brandel BR-188 density gradient fractionation system. ^35^S-Met/Cys labelling was performed as described in^36^.

## Acknowledgments

We are extremely thankful to all the labs, teams and shared resources at Janelia for supporting our research: James Zhe Liu and Peng Dong for helping us generating the knock-in mouse Embryonic Stem Cells using Crispr-Cas9. Kimberly Ritola and Hyun Ah Yi (Viral Tools) for producing lentiviral particles used in this study, Kevin McGowan (Molecular Biology) for analyzing the stability of the ARC mRNA, Kathy Schaefer and Jenny Hagemeier (Cell and Tissue Culture) for their help with single-cell sorting. We are grateful to Jim Cox and Kendra Morris (Vivarium) mouse colony management.

This work was funded by the Howard Hughes Medical Institute. MJ is supported by grants from the Canadian Institute for Health Research (CIHR) and Natural Sciences and Engineering Council (NSERC), and a salary award from the Fonds de Recherche du Québec en Santé (FRQS).

## Author Contribution

V.G. and B.P.E. performed experiments and analyzed the data. B.P.E acquired single-molecule and spectroscopy data. V.G. and M.F. generated cell lines. M.F. optimized the protocol for neuronal differentiation of mESCs. D.W. isolated the primary neuronal cultures. LP.L. performed polysome profiling and ^35^S-Met/Cys-labelling. S.P. improved algorithms for bead-based registration of imagery. V.G., B.P.E, M.J. and R.H.S wrote the manuscript.

