## Supplemental Figures for "Cap-dependent translation initiation monitored in living cells"

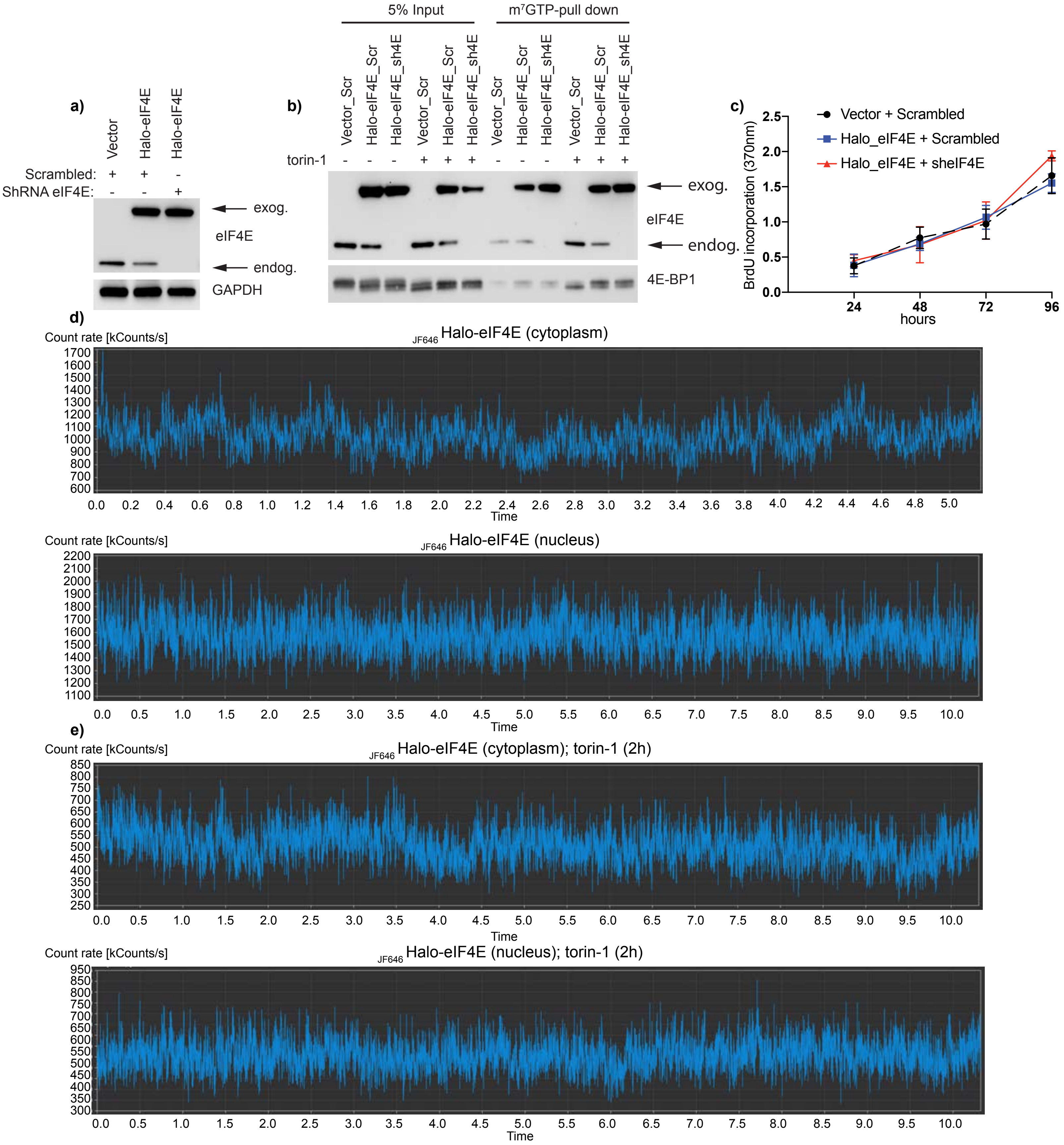


**Suppl. Fig.1 Halo-eIF4E rescue cell proliferation and binds the 5'cap mRNA.**
**a)** Vector control or Halo-eIF4E expressed in NIH3T3 cells infected with scrambled shRNA control (Scrambled) or shRNA targeting eIF4E (shRNA eIF4E). Total cell lysates were analyzed by western blotting. eIF4E antibodies detect both endogenous and exogenous eIF4E as indicated by the arrows. GAPDH was used as loading control. **b)** Cells described in a) were treated with DMSO (vehicle) or torin-1 for 1 hour. Total cell lysates (Input) were subjected to m$^7$GTP-pull down assay and analyzed with the indicated antibodies. The Halo tag does not prevent eIF4E binding to the 5'cap analogue or 4E-BP1 binding upon mTOR inhibition. **c)** Proliferation of cells described in a) was measured by 5-bromo-2′-deoxyuridine (BrdU) incorporation. The results are represented as mean absorbance at 370 nm ± s.d. from three independent experiments. **d,e)** NIH3T3 cells that express Halo-eIF4E, in which the endogenous counterpart was knocked down by shRNA, were treated with DMSO (control) or 250nM torin-1 for 2 hours. Representative examples of $_{JF646}$Halo-eIF4E fluorescent fluctuations throughout the focal volume, in the cytoplasm and in the nucleus and in the indicated conditions. Major fluctuations are only detected in the cytoplasm of translating cells.

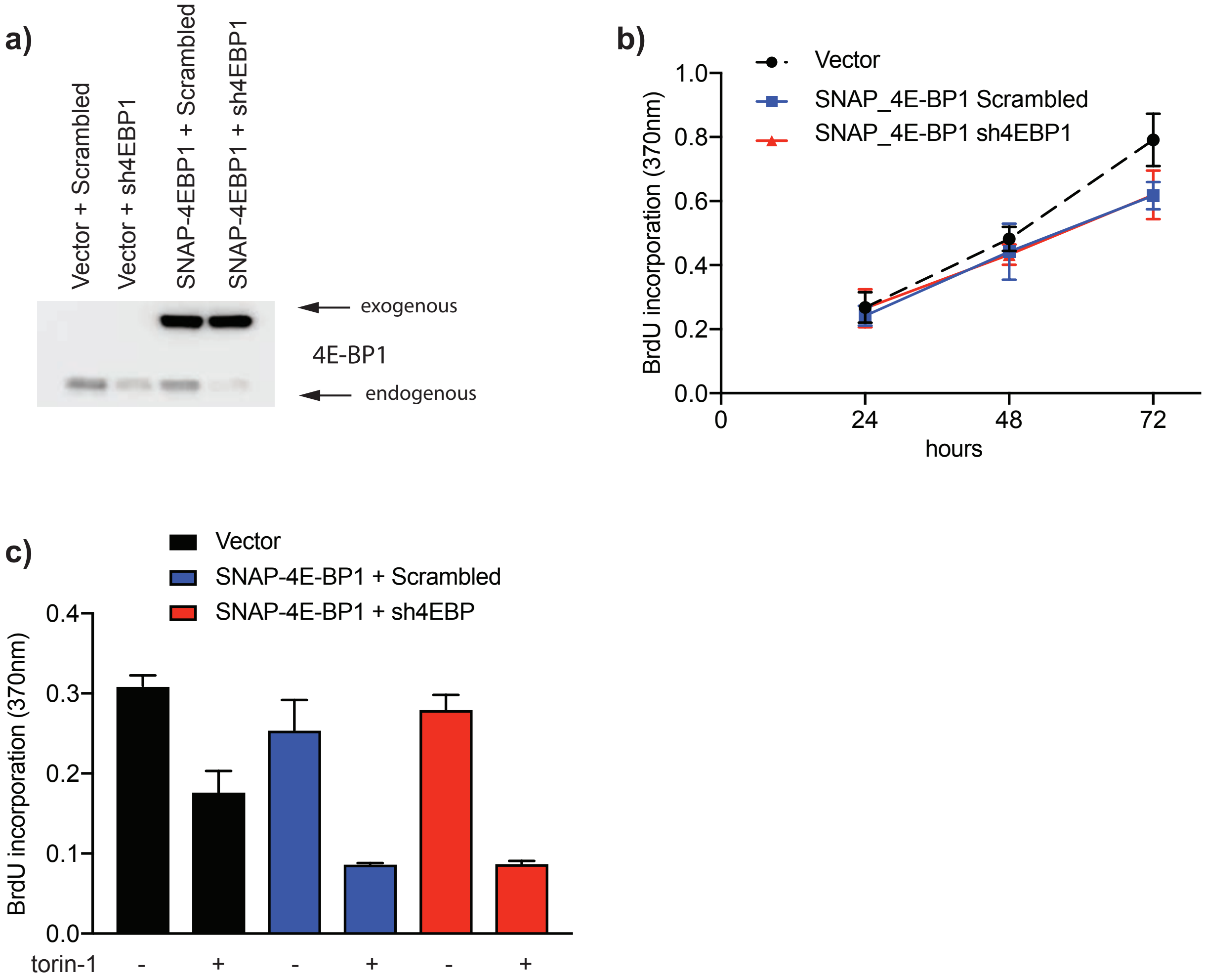


**Suppl. Fig. 2 Overexpression SNAPf-4E-BP1 slowed down cell proliferation and increased torin-1 efficacy.**
**a)** Vector control or SNAP$_f$-4E-BP1 were expressed in NIH3T3 cells infected with scrambled shRNA control (Scrambled) or shRNA targeting 4EBP1 (sh4EBP1). Total cell lysates were analyzed by western blotting. 4E-BP1 antibodies detect both endogenous and exogenous 4E-BP1 as indicated by the arrows. **b)** Proliferation of NIH3T3 cells infected with scrambled shRNA control (Scrambled) or shRNA targeting endogenous 4EBP1 (sh4EBP1) that express Vector control or SNAPf-4E-BP1 was measured by 5-bromo-2′-deoxyuridine (BrdU) incorporation. The results are represented as mean absorbance at 370 nm ± s.d. from three independent experiments. SNAPf-4E-BP1 overexpression slightly decreased cell proliferation at 48 and 72 hours. **c)** Cells described in a) were treated with vehicle control (DMSO) or 250nM torin-1 for 16 hours. Proliferation was measured as in b). The results are represented as mean absorbance at 370 nm ± s.d. from three independent experiments. 4E-BP1 overexpression increase the cytostatic effect of torin-1.

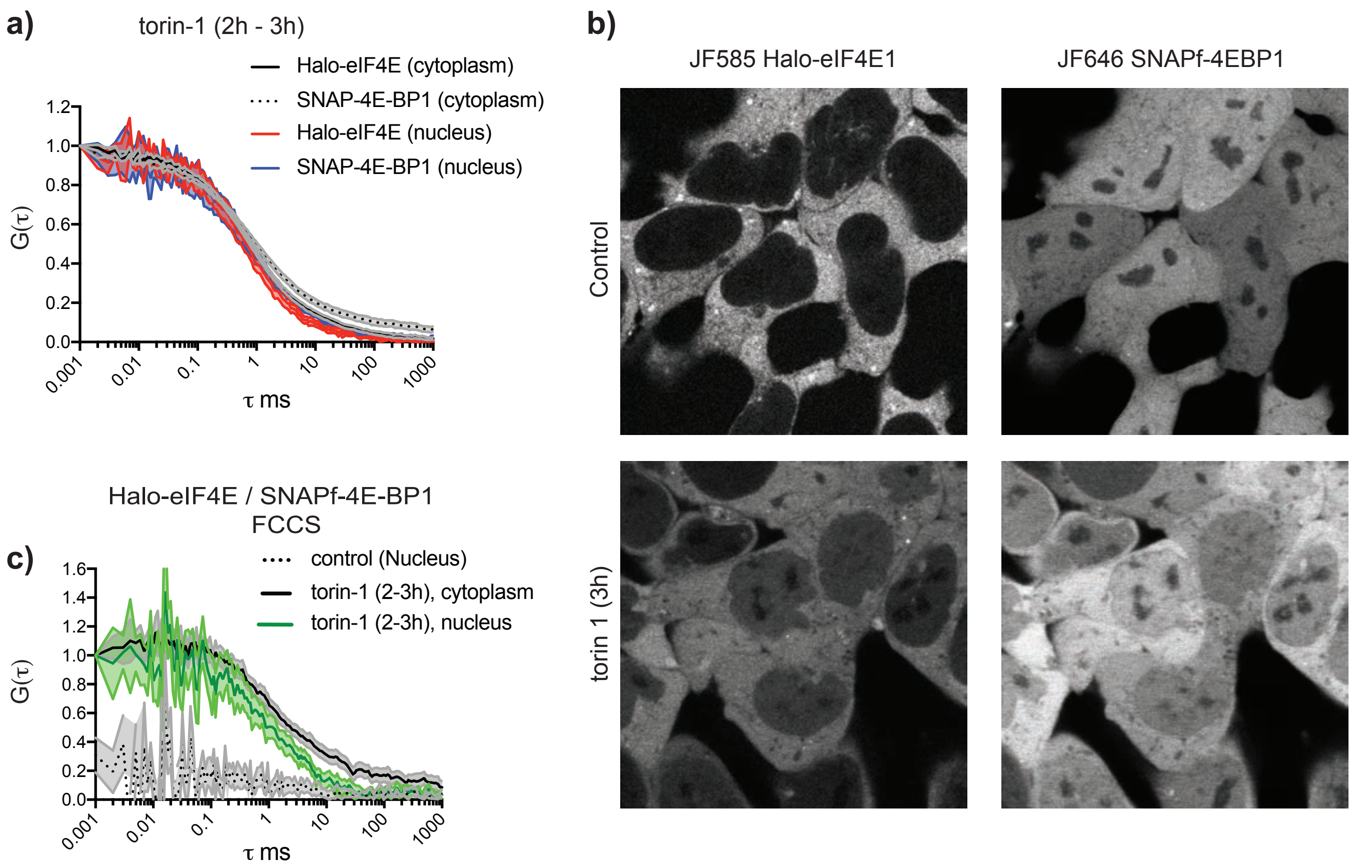


**Suppl. Fig.3 eIF4E:4E-BP1 complexes accumulate in the nucleus upon prolonged mTOR inhibition.**
**a-c)** Differentiated mESC in which Halo and $SNAP_f$ tag were inserted into the EIF4E1 and 4EBP1 locus, respectively, were treated with vehicle (DMSO) or 250nM torin-1 from 2 to 3 hours. Halo-eIF4E and $Snap_f$-4E-BP1 autocorrelations diffuse as fast as the nuclear counterparts in the indicated conditions (a). Representative field of view of differentiated mESC showing subcellular distribution of $_{JF585}$Halo-eIF4E and $_{JF646}$Snapf-4E-BP1 in control and torin-1 treated cells. Images depicted distribution of initiation factors in living cells (b). eIF4E translocate to the nucleus 3 hours upon mTOR inhibition. Simultaneous diffusion of $_{JF585}$Halo-eIF4E and $_{JF646}$Snapf-4E-BP1 analyzed in differentiated mESC by fluorescent cross-correlation spectroscopy (FCCS) in the indicated conditions. Cross-correlation was detected in the cytoplasm and in the nucleus 3 hours upon mTOR inhibition. No cross-correlation was observed in the nucleus of translating cells (control) (c).

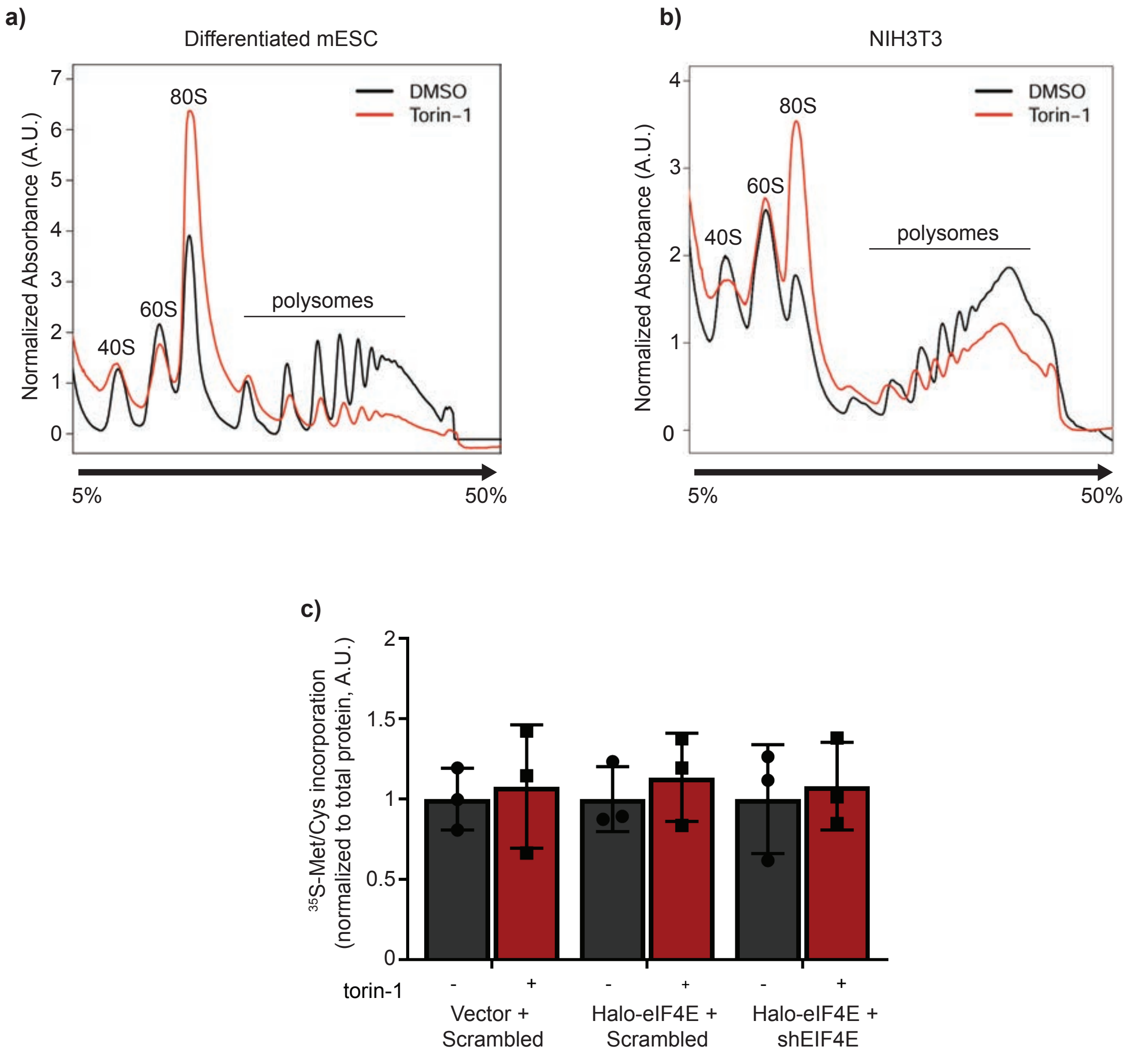


**Suppl. Fig. 4 Impaired translation initiation due to eIF4E release from the 5'cap with no major difference in global translation. a,b)** Differentiated mESC Halo-eIF4E$^{+/+}$ (a) and NIH3T3 (b) that express exogenous Halo-eIF4E were treated with DMSO (control) or 250nM torin-1 for 2 hours. Cytosolic extracts were sedimented by centrifugation on 5-50% sucrose gradients. Free ribosomal subunits (40S and 60S), monosomes (80S) and translating ribosomes (polysome) are indicated. Increased in the 80S pick and decreased polysome levels showed initiation defects upon mTOR inhibition. **c)** Global protein synthesis measure by $^{35}$S-Met/Cys incorporation in the indicated cell lines treated with DMSO (-) or torin-1 (+) for 2 hours. $^{35}$S-Met/Cys incorporation was normalized by total proteins and expressed as arbitrary units (A.U). Each replicate is represented as a single point on the corresponding bar graph. Expression of Halo-eIF4E does not sensitize the cells to torin-1 treatment.

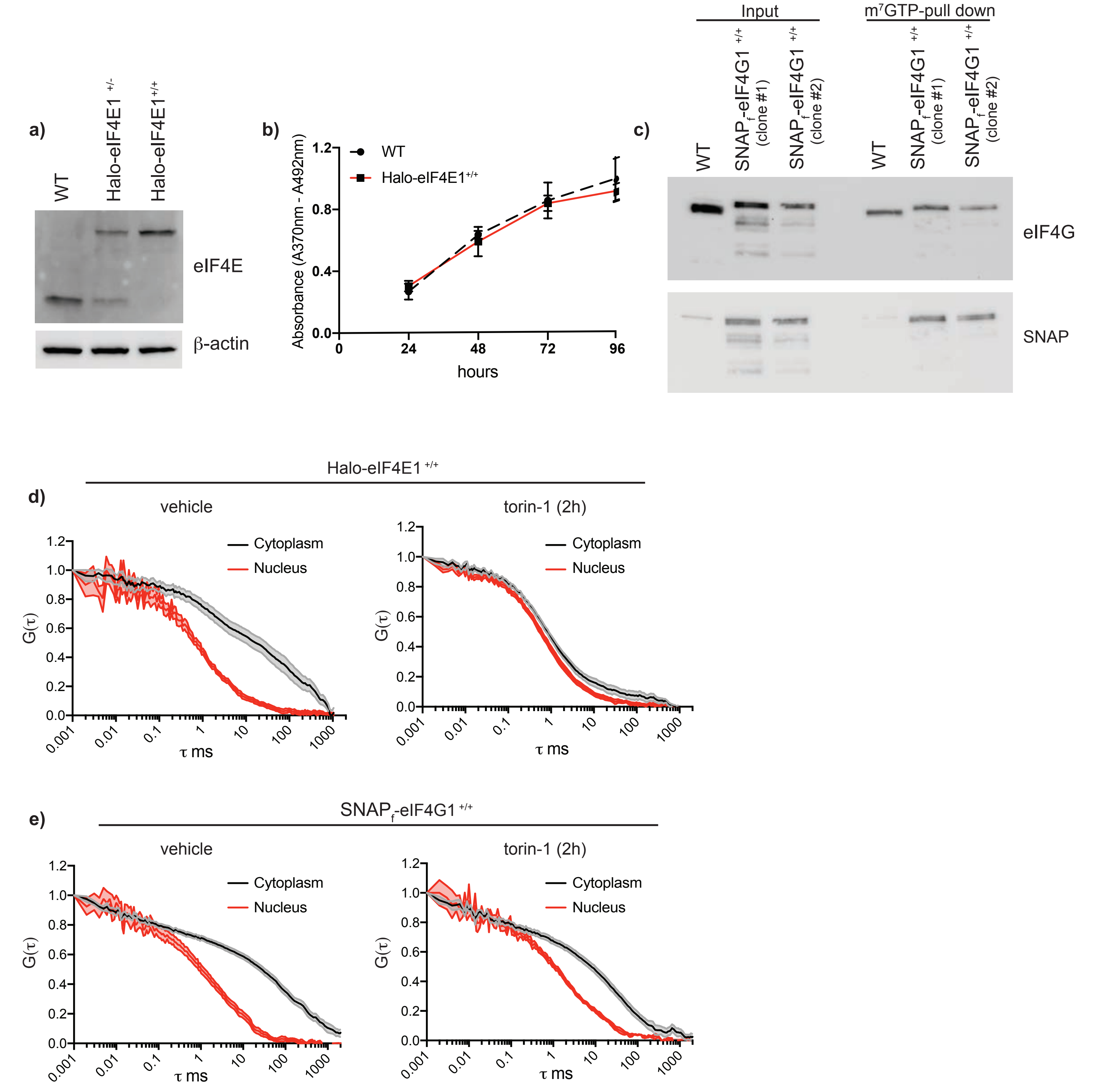


**Suppl. Fig. 5 Tagging of endogenous translation factors does not affect mESC viability or their binding dynamics.**
**a)** Halo-tag inserted in the endogenous locus of EIF4E by Crispr/cas9. Total cell lysates from parental, heterozygote and homozygote cells was analyzed by western blotting using eIF4E antibodies. β- actin was used as a loading control. Heterozygote (Halo-eIF4E$^{+/-}$) or homozygote insertion (Halo-eIF4E$^{+/+}$) is shown. **b)** Proliferation of parental and Halo-eIF4E homozygote (Halo-eIF4E$^{+/+}$) differentiated to fibroblasts was measured by 5-bromo-2′-deoxyuridine (BrdU) incorporation. The results are represented as mean absorbance at 370 nm ± s.d. from three independent experiments. **c)** SNAP$_f$-tag was inserted in the endogenous locus of EIF4G1. Total cell lysates (input) from parental and eIF4G1 homozygote differentiated mESC (SNAP$_f$-eIF4G1$^{+/+}$) were subjected to m$^7$GTP-pull down assay and analyzed by western blotting using the indicated antibodies.
**d,e)** Autocorrelation of endogenous $_{JF646}$Halo-eIF4E1 and $_{JF646}$SNAPf-eIF4G1 in cells described above. Cells were treated with DMSO (control) or 250nM torin-1 for 2 hours (2h). eIF4E and eIF4G1 molecules diffuse slower in the cytoplasmic of translating cells as compared to nuclear counterpart. Upon torin-1 treatment (torin-1 (2h)), cytoplasmic eIF4E molecules, but not eIF4G, diffuse as fast as the nuclear counterpart.

**a)**

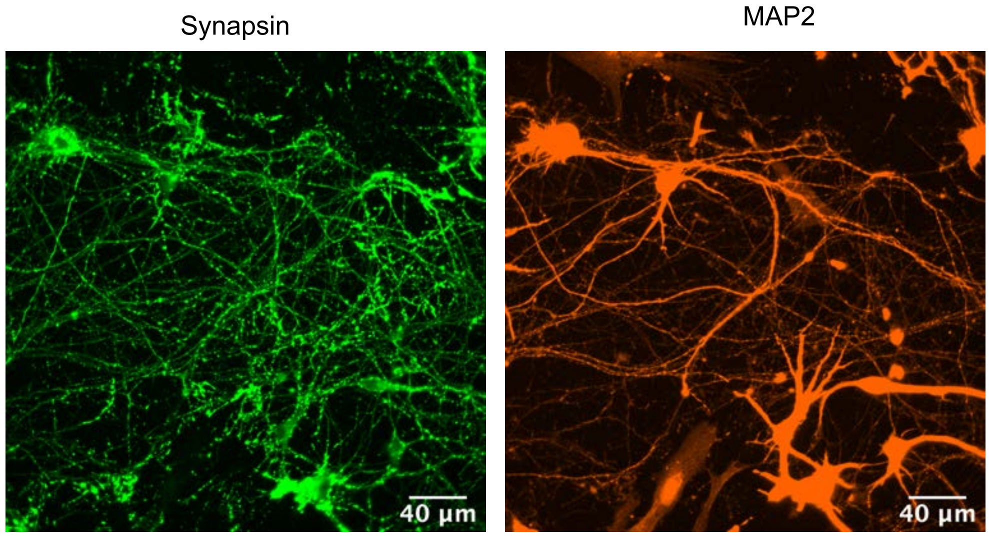


**b)**

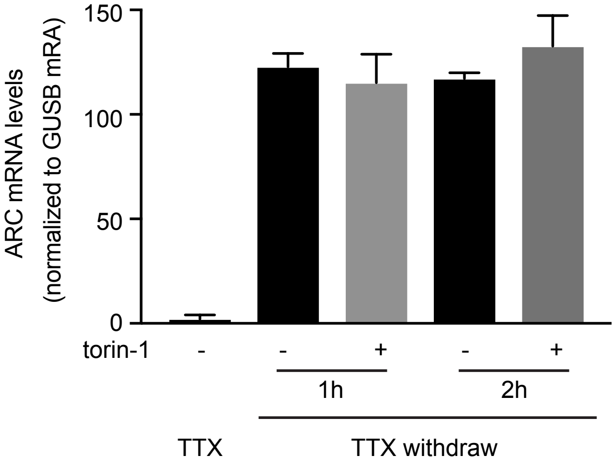


**Suppl. Fig. 6 mTOR inhibition does not affect ARC mRNA stability.**
a) Differentiated neurons derived from mESC. Immunostaining with the pre-synaptic marker Synapsin and the neuronal marker MAP2 is shown. **b)** Neurons were activated by TTX withdrawal with and without 250nM torin-1 for 1 and 2 hours. Expression levels of the mature ARC mRNA, in the indicated conditions, were determined by RT-qPCR. Values were normalized to the levels of the h ouse-keeping GUSB mRNA.
